# Changes in global brain connectivity in LSD-induced altered states of consciousness are attributable to the 5-HT2A receptor

**DOI:** 10.1101/219956

**Authors:** Katrin H. Preller, Joshua B. Burt, Jie Lisa Ji, Charles Schleifer, Brendan Adkinson, Philipp Stämpfli, Grega Repovs, John H. Krystal, John D. Murray, Franz X. Vollenweider, Alan Anticevic

## Abstract

Lysergic acid diethylamide (LSD) is a psychedelic drug with predominantly agonist activity at various serotonin (5-HT) and dopamine receptors. Despite the therapeutic and scientific interest in LSD, the specific receptor contributions to its neurobiological effects remain largely unknown. To address this knowledge gap, we conducted a double-blind, randomized, counterbalanced, cross-over study during which 24 healthy participants received either i) placebo+placebo, ii) placebo+LSD (100 μg po), or iii) ketanserin – a selective 5-HT2A receptor antagonist. Here we focus on resting-state fMRI, a measure of spontaneous neural fluctuations that can map functional brain connectivity. We collected resting-state data 75 and 300 minutes after LSD/placebo administration. We quantified resting-state functional connectivity via a fully data-driven global brain connectivity (GBC) method to comprehensively map LSD neuropharmacological effects. LSD administration caused widespread GBC alterations that followed a specific topography: LSD reduced connectivity in associative areas, but concurrently increased connectivity across sensory and somatomotor areas. The 5-HT2A receptor antagonist, ketanserin, fully blocked the subjective and neural LSD effects. We show that whole-brain data-driven spatial patterns of LSD effects matched 5-HT2A receptor cortical gene expression in humans, which along with ketanserin effects, strongly implicates the 5-HT2A receptor in LSD’s neuropharmacology. Critically, the LSD-induced subjective effects were associated with somatomotor networks GBC changes. These data-driven neuropharmacological results pinpoint the critical role of 5-HT2A in LSD’s mechanism, which informs its neurobiology and guides rational development of psychedelic-based therapeutics

## Significance Statement

Lysergic acid diethylamide (LSD) is a psychedelic drug. Interest in psychedelic neurobiology is reinforced by their safe use for research and clinical applications. However, there are major knowledge gaps regarding LSD’s neuropharmacology. Using cutting-edge neuroimaging methods we address knowledge gaps in LSD’s neuropharmacology by comprehensively mapping LSD’s effect on data-driven neural connectivity along with its reversal via specific serotonin receptor blockade. For the first time, we report a disintegration of executive and integration of sensory-motor networks induced by LSD that is predominantly dependent on serotonin 2A receptor activation. Furthermore, we implicate the somatomotor network brain-wide integration in LSD-induced subjective effects. These results inform the neurobiology of altered states of consciousness, with critical implications for rational development of novel psychedelic-based therapeutics.

## INTRODUCTION

Disorders of perception and the form and content of thought are important contributors to the global burden of disease (1). Mechanistic studies of consciousness may be undertaken using psychedelic drugs as pharmacologic probes of molecular signaling within cortical networks underlying perception and thought. In particular, lysergic acid diethylamide (LSD) is a psychedelic drug with predominantly agonist activity at serotonin (5-HT)-2A/C, -1A/B, -6, and -7 and dopamine D2 and D1 receptors (R). Its administration produces characteristic alterations in perception, mood, thought, and the sense of self (2–4). Despite its powerful effects on consciousness, human research on LSD neurobiology stalled in the late 1960s because of a narrow focus on the experiential effects of hallucinogenic drugs, combined with a lack of understanding of its effects on molecular signaling mechanisms in the brain. However, renewed interest in the potentially beneficial clinical effects of psychedelics (5–8) warrants a better understanding of their underlying neuropharmacology. Nevertheless, major knowledge gaps remain regarding LSD’s neurobiology in humans as well as its time-dependent receptor neuropharmacology.

To address this critical gap, the current study aims to comprehensively map time-dependent pharmacological effects of LSD on neural functional connectivity in healthy human adults. The goal is to leverage the statistical properties of the slow (<1 Hz) intrinsic fluctuations of the blood-oxygen-level-dependent (BOLD) signal hemodynamics at rest (i.e. resting-state functional connectivity (rs-fcMRI)). Critically, rs-fcMRI analyses are able to reveal the functional architecture of the brain, which is organized into large-scale systems exhibiting functional relationships across space and time (9–11). Rs-fcMRI measures have furthermore revealed potential biomarkers of various neural disorders (12–14), as well as proven sensitive to the effects of neuropharmacological agents (15, 16).

Focused analyses on specific regions revealed effects of intravenously administered LSD on functional connectivity between V1 and distributed cortical and subcortical regions (17). However, such ‘seed-based’ approaches rely on explicitly selecting specific regions of interest based on a priori hypotheses. Therefore, such an approach has limited ability to detect pharmacologically-induced dysconnectivity not predicted *a priori*. To characterize LSD effects on functional connectivity in the absence of strong a priori hypotheses, the current study employed a fully data-driven approach derived from graph theory called Global Brain Connectivity (GBC) (18). In essence, GBC computes the connectivity of every voxel in the brain with all other voxels and summarizes that in a single value. Therefore, areas of high GBC are highly functionally connected with other areas and might play a role in coordinating large-scale patterns of brain activity (19). Reductions in GBC may indicate decreased participation of a brain area in larger networks, whereas increased GBC may indicate a broadening or synchronization of functional networks (18). One small study examined GBC after intravenously administered LSD in a sample of 15 participants, revealing connectivity elevations across higher-order association cortices (20). While compelling, this preliminary connectivity study did not take into account the influence of global signal (GS) artifacts (e.g. via global signal regression, GSR), which are known to exhibit massive differences in clinical populations and pharmacological manipulations (21–24). Specifically, GS is hypothesized to contain large amounts of non-neuronal artifact (e.g., physiological, movement, scanner-related) (25), which can induce spuriously high correlations across the brain (11). No data-driven study has comprehensively characterized LSD-induced changes as a function of rigorous artifact removal. Thus, a major objective here was to map data-driven LSD-induced dysconnectivity in the context of GS removal, which is key for separating signal and noise.

Another aim of the current study was to determine the extent to which the neural and behavioral effects of LSD are mediated by 5HT2A receptors. Preclinical studies suggest that LSD binds potently to many neuroreceptors including 5HT2A, 5HT2C, 5HT1A, D2, and other receptors (2, 26). Yet, a recent paper from our group (27) reported that the psychedelic effects of LSD were entirely blocked in humans by ketanserin, a selective antagonist at 5-HT2A and α-adreno receptors (28). This would suggest that the neural effects of LSD should be blocked by ketanserin. It also suggests that networks modulated by LSD should highly associated with the distribution of 5HT2A receptors in the brain and not closely associated with the distribution of receptors unrelated to the mechanism of action of LSD.

Here we leverage recent advances (29) in human cortical gene expression mapping to inform the spatial topography of neuropharmacologically-induced changes in data-driven connectivity. We hypothesized that the LSD-induced GBC change will quantitatively match the spatial expression profile of genes coding for the 5-HT2A receptor. In turn, we hypothesized that this effect will be preferential for the 5-HT2A but not other receptors and that the spatial match will be vastly improved after artifact removal. In doing so, this convergence of neuropharmacology and gene expression mapping validates the contribution of the 5-HT2A receptor to LSD neuropharmacology. In turn, it also highlights a general method for relating spatial gene expression profiles to neuropharmacological manipulations, which has direct and important implications for the rational refinement of any receptor neuropharmacology.

Collectively, this pharmacological neuroimaging study addresses the following major knowledge gaps in our understanding of LSD neurobiology, by demonstrating: i) data-driven LSD effects across brain-wide networks, which are exquisitely sensitive to artifact removal, ii) the subjective and neural effects of LSD neuropharmacology are attributable to the 5-HT2A receptor, and iii) the cortex-wide LSD effects can be mapped onto the spatial expression profile of the gene coding for the 5-HT2A receptor.

## RESULTS

### LSD Modulates Global Brain Connectivity and Induces Marked Subjective Drug Effects

The main effect of drug on GBC computed with GSR revealed significant (TFCE type 1 error protected, 10000 permutations) widespread differences in GBC between drug conditions in cortical and subcortical areas (**Fig. 1A**). Comparing LSD to (Ket+LSD)+Pla conditions across sessions shows that LSD induces hyper-connectivity predominately in sensory areas, i.e. the occipital cortex, the superior temporal gyrus, and the postcentral gyrus, as well as the precuneus. Hypo-connectivity was induced in subcortical areas as well as cortical areas associated with associative networks, such the medial and lateral prefrontal cortex, the cingulum, the insula, and the temporoparietal junction. All changes in connectivity were expressed bilaterally (**Fig. 1A**). **Fig. 1B** shows mean connectivity strength (Fz) for each drug condition and the distribution of Fz values within voxels showing significant hyper- and hypo-connectivity for LSD compared to (Ket+LSD)+Pla conditions. Mean Fz values do not differ between Pla and Ket+Pla conditions either in hyper-connected or in hypo-connected areas. **Fig. 1C** depicts the comparison between LSD and Pla conditions and **Fig. 1D** the comparison between LSD and Ket+LSD conditions. Similarly to the comparison LSD vs. (Ket+LSD)+Pla shown in **Fig. 1A**, LSD compared to both Pla and Ket+LSD separately induced a connectivity pattern characterized by significant (TFCE type 1 error protected, 10000 permutations) hyper-connectivity in predominantly sensory areas and significant hypo-connectivity in associative networks. The similarity between the LSD>Pla and LSD>Ket+LSD contrasts is corroborated by a significant positive correlation (r=0.91, p<0.001) between the respective Z-maps (**Fig. 1E**). Furthermore, only negligible differences were observed when comparing Ket+LSD and placebo conditions directly (**Fig. S3**). Together, these results indicate that LSD-induced GBC alterations are predominantly attributable to its agonistic activity onto the 5-HT2A receptor. In line with this, a repeated-measures ANOVA (drug condition × scale) was conducted for the retrospectively administered Altered States of Consciousness (5D-ASC) questionnaire, and revealed significant main effects for treatment (F(2, 46) = 88.49, p <0.001) and scale (F(10, 230) = 14.47, p<0.001), and a significant interaction of treatment × scale (F(20, 460) = 13.02, p<0.001). Bonferroni corrected simple main effect analyses showed increased ratings on all 5D-ASC scales in the LSD condition compared to Pla and Ket+LSD conditions (all p<0.05) except for the scales spiritual experience and anxiety (all p>0.20). Pla and LSD+Ket scores did not differ on any scale (all p > 0.90) (**Fig. 1F**).

**Figure 1.**
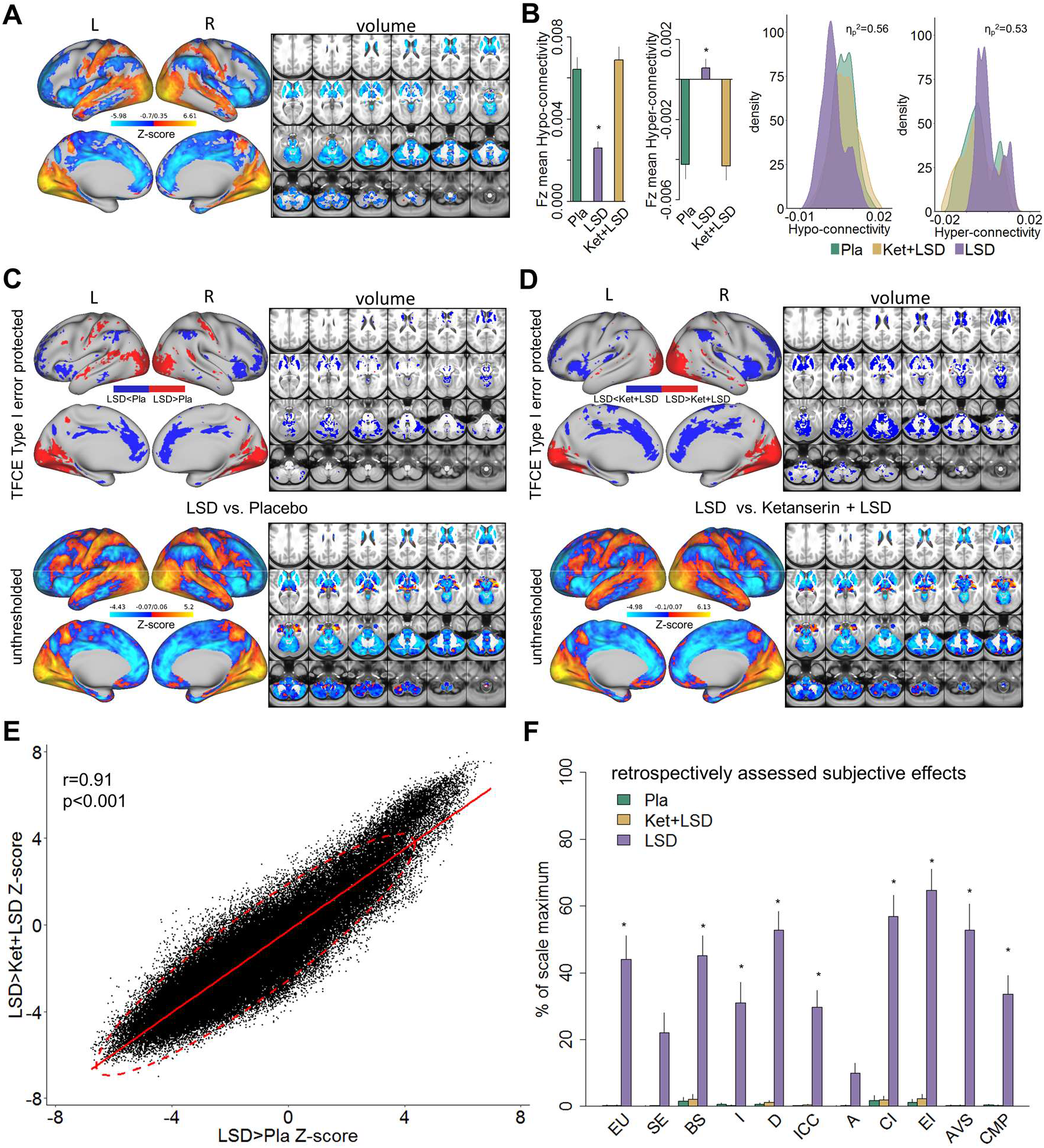
Effect of drug condition on Global Brain Connectivity and subjective drug effects. **(A)** Z-score map for the effect of LSD condition vs. Ketanserin+LSD condition + Placebo condition within areas showing a significant main effect of drug (TFCE type 1 error protected). Red/orange areas indicate regions where participants exhibited stronger GBC in the LDS condition, whereas blue areas indicate regions where participants exhibited reduced GBC condition, compared with (Ketanserin+LSD)+Placebo conditions. **(B)** Bar plots show mean connectivity strength (Fz) values for hyper- and hypo-connected areas averaged across voxels showing a significant main effect of drug. Distribution plots show distribution of connectivity strength (Fz) values within voxels showing significant hyper- and hypo-connectivity for LSD compared to (Ketanserin+LSD)+Placebo conditions. **(C)** Top panel displays significant (TFCE type 1 error protected) areas showing increased (red) and decreased (blue) GBC in the LSD condition compared to Placebo. Lower panel shows the corresponding unthresholded Z-score map. Red/orange areas indicate regions where participants exhibited stronger GBC in the LSD condition, whereas blue areas indicate regions where participants exhibited reduced GBC in the LSD condition, compared with Placebo condition. **(D)** Top panel displays significant (TFCE type 1 error protected) areas showing increased (red) and decreased (blue) GBC in the LSD condition compared to Ketanserin+LSD. Lower panel shows the corresponding unthresholded Z-score map. Red/orange areas indicate regions where participants exhibited stronger GBC in the LSD condition, whereas blue areas indicate regions where participants exhibited reduced GBC in the LSD condition, compared with Ketanserin+LSD condition. **(E)** Scatterplot showing a positive relationship between drug condition differences in GBC. Plotted are Z-scores for all voxels for the LSD>Placebo comparison (see panel C, X-axis) and LSD>Ketanserin+LSD comparison (see panel D, Y-axis). Ellipse marks the 95% confidence interval. **(F)** Retrospectively assessed (720 min after second drug administration) subjective drug-induced effects. Effects were assessed with the Five Dimension Altered States of Consciousness Questionnaire. EU: Experience of Unity; SE: Spiritual Experience; BS: Blissful State; I: Insightfulness; D: Disembodiment; ICC: Impaired Control and Cognition; A: Anxiety; CI: Complex Imagery; EI: Elementary Imagery; AVS: Audio-Visual Synesthesia; CMP: Changed Meaning of Percepts. N=24. * indicates significant difference between LSD and Pla, and LSD and Ketanserin+LSD drug conditions, p<0.05, Bonferroni corrected.

### Significant Relationship Between Hyper- and Hypo-connectivity

To test whether the directionality of LSD-induced effects on GBC (hyper-connectivity across sensory networks, hypo-connectivity across associative networks) are separable effects or result from functionally related systems-level perturbations, we correlated the mean connectivity strength difference between placebo and LSD in hyper-connected regions with mean connectivity strength difference in hypo-connected areas (based on the LSD vs (Ketanserin+LSD)+Placebo contrast) across subjects. There was a significant correlation between hypo- and hyper-connectivity (r=-0.90, p<0.001, **Fig. 2**) indicating that participants with the highest LSD-induced coupling within sensory networks also showed the strongest LSD-induced decoupling in associative networks. This suggests that LSD-induced alterations in information flow across these networks may result from systems-level perturbations.

**Figure 2.**
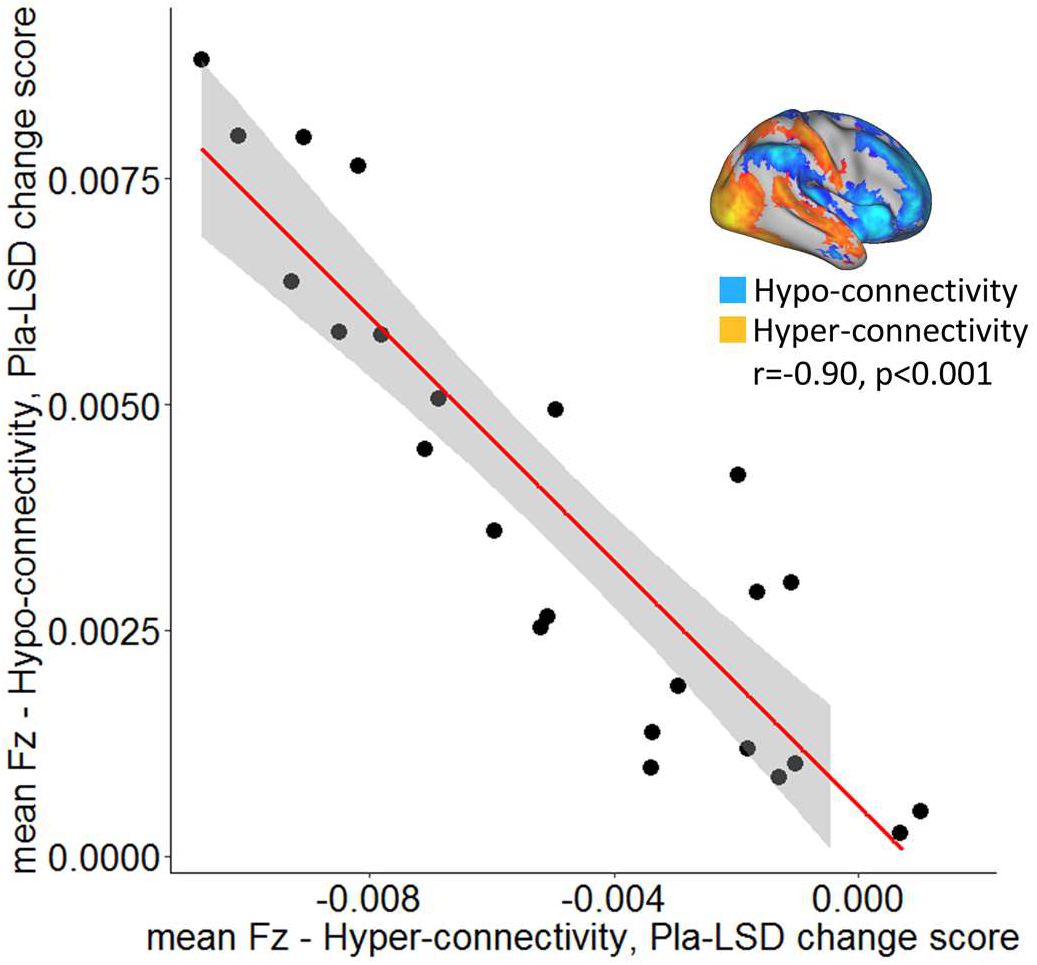
Relationship between hyper- and hypo-connectivity across subjects. Scatterplot showing significant negative relationship evident between averaged hyper- and hypo- connected voxels (based on the LSD vs (Ketanserin+LSD)+Placebo contrast, see Fig. 1A and inlet) across subjects (black data points) for placebo – LSD condition change scores. Grey background indicates the 95% confidence interval. N=24.

### Time Course of Subjective Drug Effects

To investigate the time course of subjective effects, a short version of the 5D-ASC was administered 180 min, 250 min, and 360 min after the second drug administration. A repeated-measures (treatment × time × scale) ANOVA for the short-version 5D-ASC questionnaire revealed significant main effects for treatment (F(2, 44) = 58.32, p<0.001), time (F(2, 44) = 26.61, <0.001), and scale (F(4, 88) = 14.83, p<0.001) and significant interactions for treatment × time (F(4, 88) = 16.89, p<0.001), treatment × scale (F(8, 176) = 12.82, p<0.001), time × scale (F(8, 176) = 4.05, p<0.001), and treatment × time × scale (F(16, 352) = 2.22, p<0.01). Bonferroni-corrected simple main effect analyses revealed that score in the LSD treatment condition differed significantly from score in the Pla and Ket+LSD treatment conditions for the blissful state scale, disembodiment scale, elementary imagery scale, and changed meaning of percepts scale at 180 and 250 min after treatment intake (all p<0.05). 360 min after intake, score on the disembodiment scale and elementary imagery scale was significantly greater in the LSD treatment condition than in the Pla and Ket+LSD treatment conditions. Scores did not differ between the Pla and Ket+LSD treatment conditions for any scale at any time point (all p > 0.90; **Fig. 3**).

**Figure 3.**
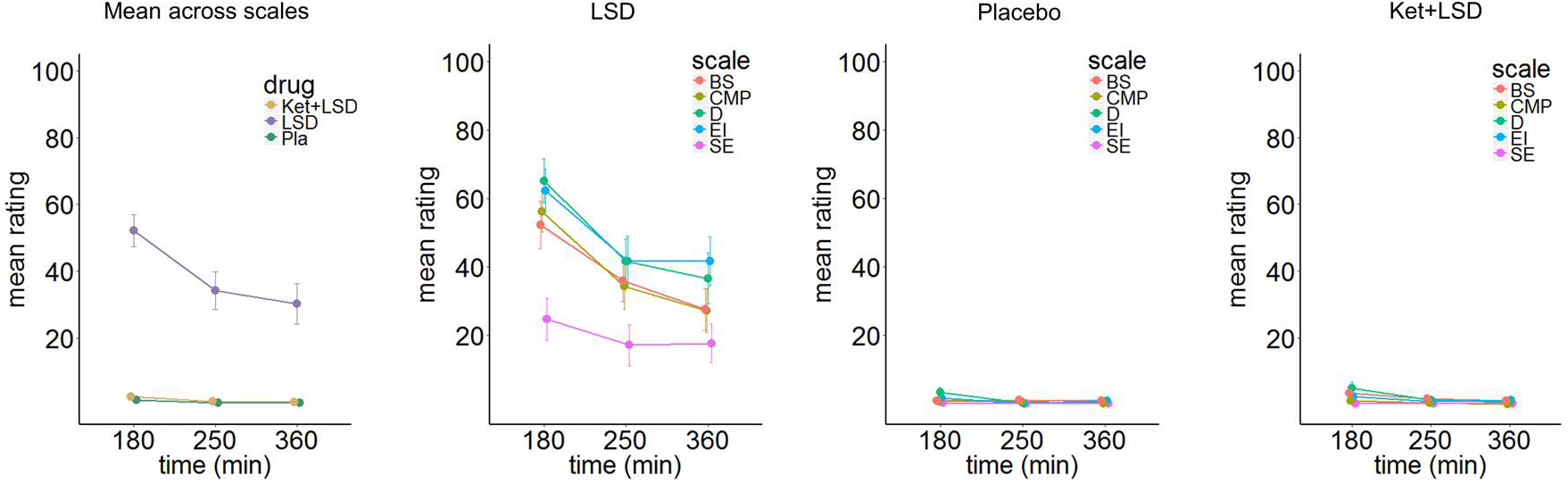
Time course of subjective drug effects. Five Dimension Altered States of Consciousness Questionnaire short version scores assessed at 180, 250, and 300 minutes after second drug administration for the means across scales, and scale scores for Placebo, LSD , and Ketanserin+LSD conditions. Scores are expressed as percent of the scale maximum. Data are expressed as means ± the standard error of the mean (SEM). BS: Blissful State; CMP: Changed Meaning of Percepts; D: Disembodiment; EI: Elementary Imagery; SE: Spiritual Experience. N= 23.

### Session affects Global Brain Connectivity in Ketanserin+LSD condition

To investigate the potentially distinct temporal phases of LSD pharmacology (30, 31), two resting-state scans were conducted on each test day: 75 minutes (session 1) and 300 minutes (session 2) after the second drug administration. No significant differences in GBC were observed when comparing session 1 and 2 within the placebo and the LSD condition (**Fig. S4**). Within the Ketanserin+LSD condition, participants showed significant decreases in GBC in session 2 compared to session 1 predominantly in occipital areas. Increases in GBC in session 2 were found in cortical regions such as the anterior and posterior cingulate cortex, and the temporoparietal junction, as well as subcortical structures including the thalamus and the basal ganglia (**Fig. 4A**). Fig. 4B shows the significant (p< 0.05) difference in mean Fz between session 1 and session 2 in hyper- and hypo-connected areas and the distribution of Fz values for voxels within hyper- and hypo-connected areas for both sessions (hyper- and hypo-connected areas are derived from the LSD vs (Ket+LSD)+Pla contrast, see **Fig. 1A**).

**Figure 4.**
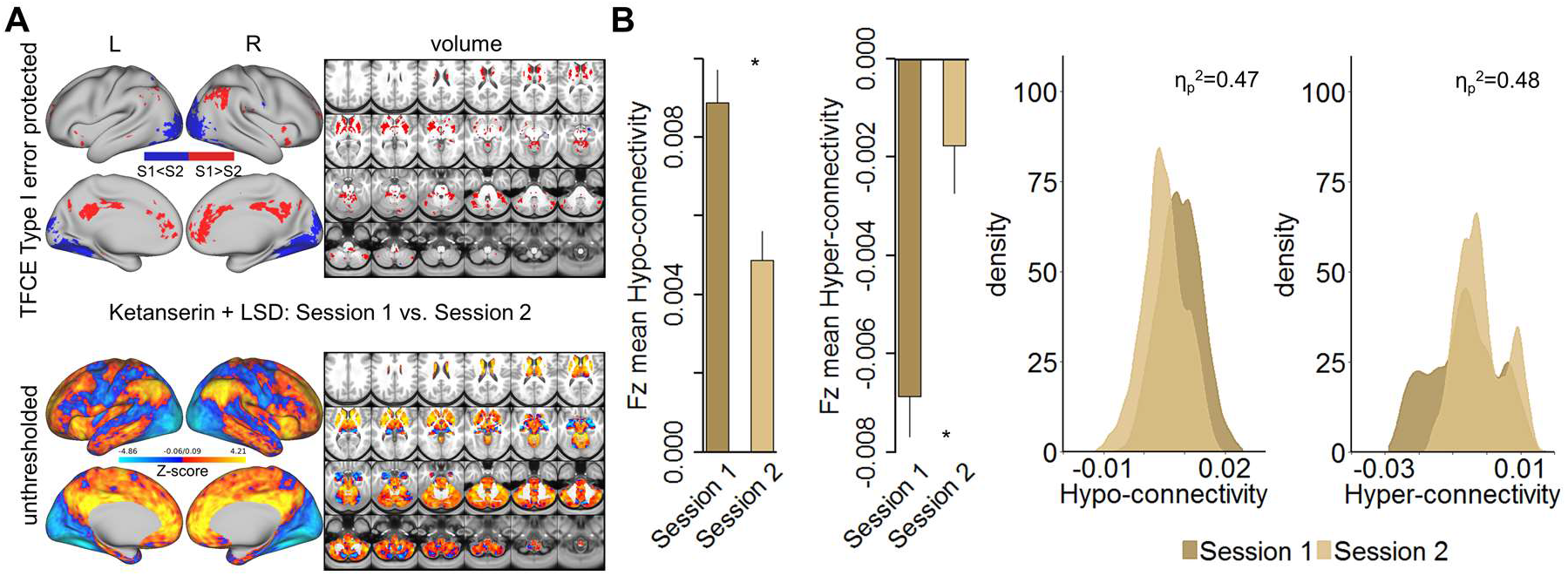
Effect of session on Global Brain Connectivity in the Ketanserin+LSD condition. **(A)** Top panel displays significant (TFCE type 1 error protected) areas showing increased (red) and decreased (blue) GBC in session 1 (75 minutes after second drug administration) compared to session 2 (300 minutes after second drug administration). Lower panel shows the corresponding unthresholded Z-score map. Red/orange areas indicate regions where participants exhibited stronger GBC in session 1, whereas blue areas indicate regions where participants exhibited reduced GBC in session 2. **(B)** Bar plots show mean connectivity strength (Fz) values for hyper- and hypo-connected areas (significant for the LSD vs (Ketanserin+LSD)+Placebo contrast) for session 1 and session 2 in the Ketanserin+LSD condition. Distribution plots show distribution of connectivity strength (Fz) values within voxels showing hyper- and hypo-connectivity (significant for the LSD vs. Ketanserin+LSD)+Placebo contrast) for session 1 and session 2 in the Ketanserin+LSD condition. * indicates significant difference between session 1 and session 2 (p<0.05). N=24.

### Influence of Global Signal Regression on Global Brain Connectivity

To investigate the influence of GSR on the results, the analysis of effect of drug condition on GBC (**Fig. 1**) was repeated without GSR. The main effect of drug on GBC computed without GSR revealed significant (TFCE type 1 error protected, 10000 permutations) predominantly left-hemispheric widespread differences in GBC between drug conditions (**Fig. S5A**). **Fig. S5B** shows mean Fz for each drug condition and the distribution of Fz values within voxels showing significant hyper- and hypo-connectivity for LSD compared to (Ket+LSD)+Pla conditions. Mean Fz values for hypo-connected voxels differed significantly between Pla and Ket+LSD conditions. Mean Fz values for hyper-connected voxels differed significantly between Pla and Ket+LSD, and LSD and Ket+LSD conditions. **Fig. S5C** depicts the comparison between LSD and Pla conditions. Without GSR LSD induced hypo-connectivity mainly in the right insula and hyper-connectivity predominantly in the cerebellum. **Fig. S5D** shows the comparison between LSD and Ket+LSD conditions with LSD-induced hypo-connectivity in the left insula and widespread predominantly left-hemispheric hyper-connectivity in the frontal and temporal cortex, the tempoparietal junction, and the cerebellum. Comparing Ket+LSD and Placebo conditions revealed Ket+LSD induced hyper-connectivity predominantly in the right hemisphere (**Fig. S5E**). LSD>Pla and LSD>Ket+LSD Z-maps were significantly correlated (r=0.81, p<0.001, **Fig. S5F**).

Given these widespread differences between GBC analysis with and without GSR, we next specifically investigated mean Fz (with and without GSR) for all drug conditions and sessions for voxels within seven functionally-defined networks using parcellations derived by Yeo et al. (32), Buckner et al. (33) and Choi et al. (34) (**Fig. 5**). This parcellation contains both sensory (visual and somatomotor) and associative (dorsal attention, ventral attention, limbic, frontoparietal control, and default mode) networks. Repeated-measures ANOVAs revealed significant main effects for drug condition for all networks (all p<0.05) except for the dorsal attention network when including GSR, with the LSD condition differing significantly from both, Pla and Ket+LSD conditions (all p<0.05, Bonferroni corrected), except for the somatomotor network, where LSD differed significantly only from Ket+LSD. Placebo and Ket+LSD conditions did not differ significantly in any network. Without GSR, main effects for drug were found in the frontoparietal control network [F(2.46)=4.09, p<0.03] with significantly lower values in the Ket+LSD condition than in both, the LSD and Pla condition, and dorsal attention network [F(2,46)=3.86, p<0.04] with significantly lower values in the Ket+LSD condition than in the Pla condition.

**Figure 5.**
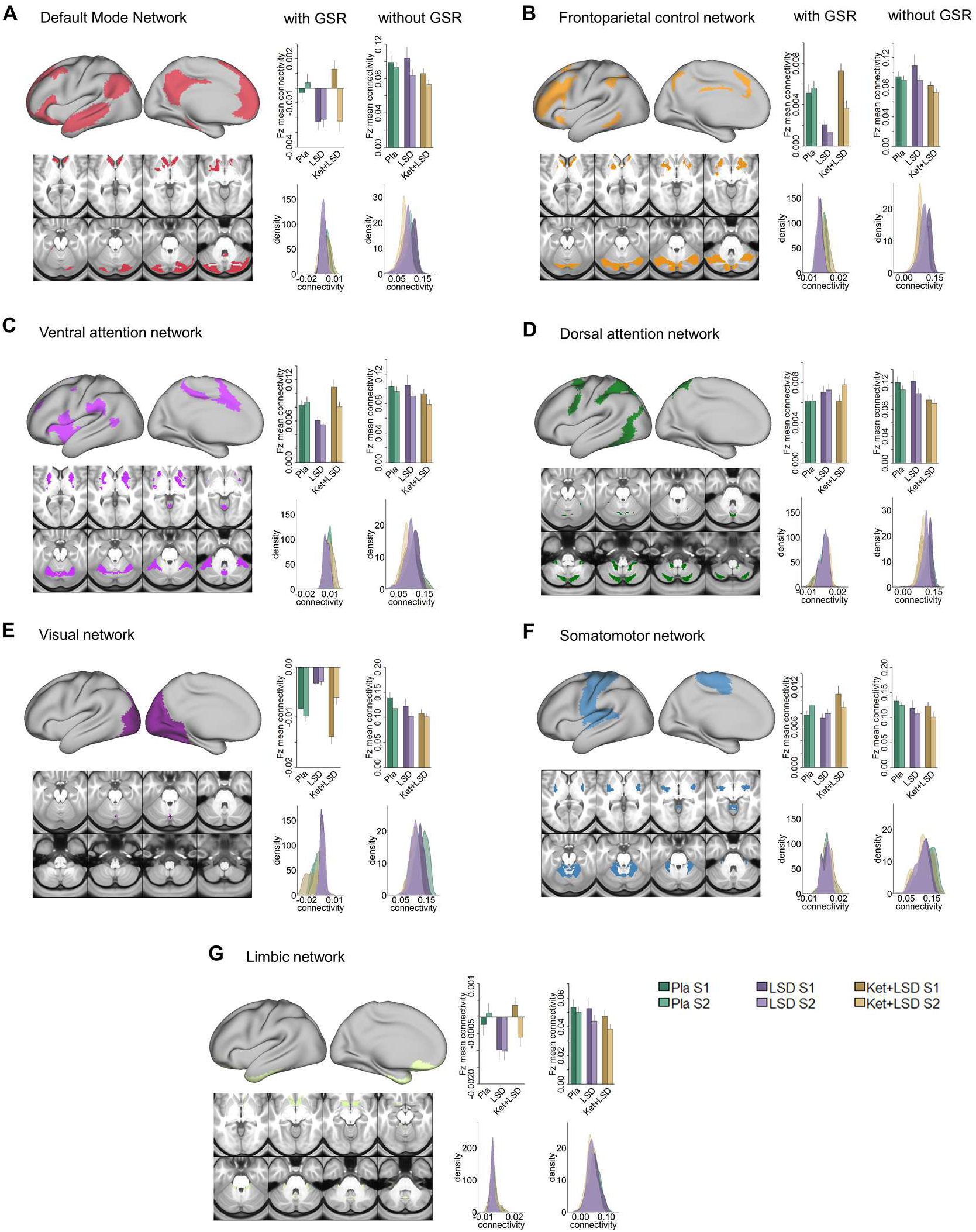
Effect of drug condition, session, and GSR on Global Brain Connectivity in functional networks (A-G). Brain maps illustrate lateral, medial, and subcortical view of functional networks. Bar plots show mean connectivity strength (Fz) values for voxels within functional networks for Placebo, LSD, and Ketanserin+LSD conditions, for session 1 and session 2 respectively, as well as with and without GSR. Distribution plots show distribution of connectivity strength (Fz) values for voxels within functional networks for Placebo, LSD, and Ketanserin+LSD conditions, for session 1 and session 2 respectively, as well as with and without GSR. N=24.

To test the relationship between hyper- and hypo-connectivity when GSR was not performed, we correlated the mean Fz difference between placebo and LSD without GSR in hyper-connected regions with the mean Fz difference in hypo-connected areas (based on the LSD vs (Ketanserin+LSD)+Placebo contrast) across subjects. There was a significant positive correlation between hyper- and hypo-connectivity (r = 0.92, p < 0.001, **Fig. S6A**). In contrast to the analysis performed with GSR showing a negative relationship between hyper- and hypo-connectivity change scores, the analysis without GSR indicates that participants with the highest LSD-induced hyperconnectivity showed the weakest LSD-induced de-coupling. Correlating the combined hyper- and hypoconnectivity values with GSR with those without GSR showed that these are not significantly related within subjects (r=0.003, p=0.99, **Fig. S6B**).

### Global Brain Connectivity in Somatomotor Network correlates with Subjective Effects

To evaluate the relationship between LSD-induced changes in GBC in functional networks and subjective LSD-induced effects, Fz mean connectivity change (LSD – Pla condition, session 2, with GSR) in the seven functional networks (see **Fig. 5**) was correlated with the mean 5-DASC short version score at 250 mins (assessment closest in time to rsfMRI data collection, see **Fig. S1** and **Fig. 3**). Correlating measures at session 2 allows high stability in LSD-induced effects. Bonferroni corrected correlations showed a significant relationship between the change in Fz connectivity in the somatomotor network and subjective LSD-induced effects (r=0.81, p<0.001, Bonferoni corrected, **Fig 6A** and **Fig. 6B**).

**Fig. 6.**
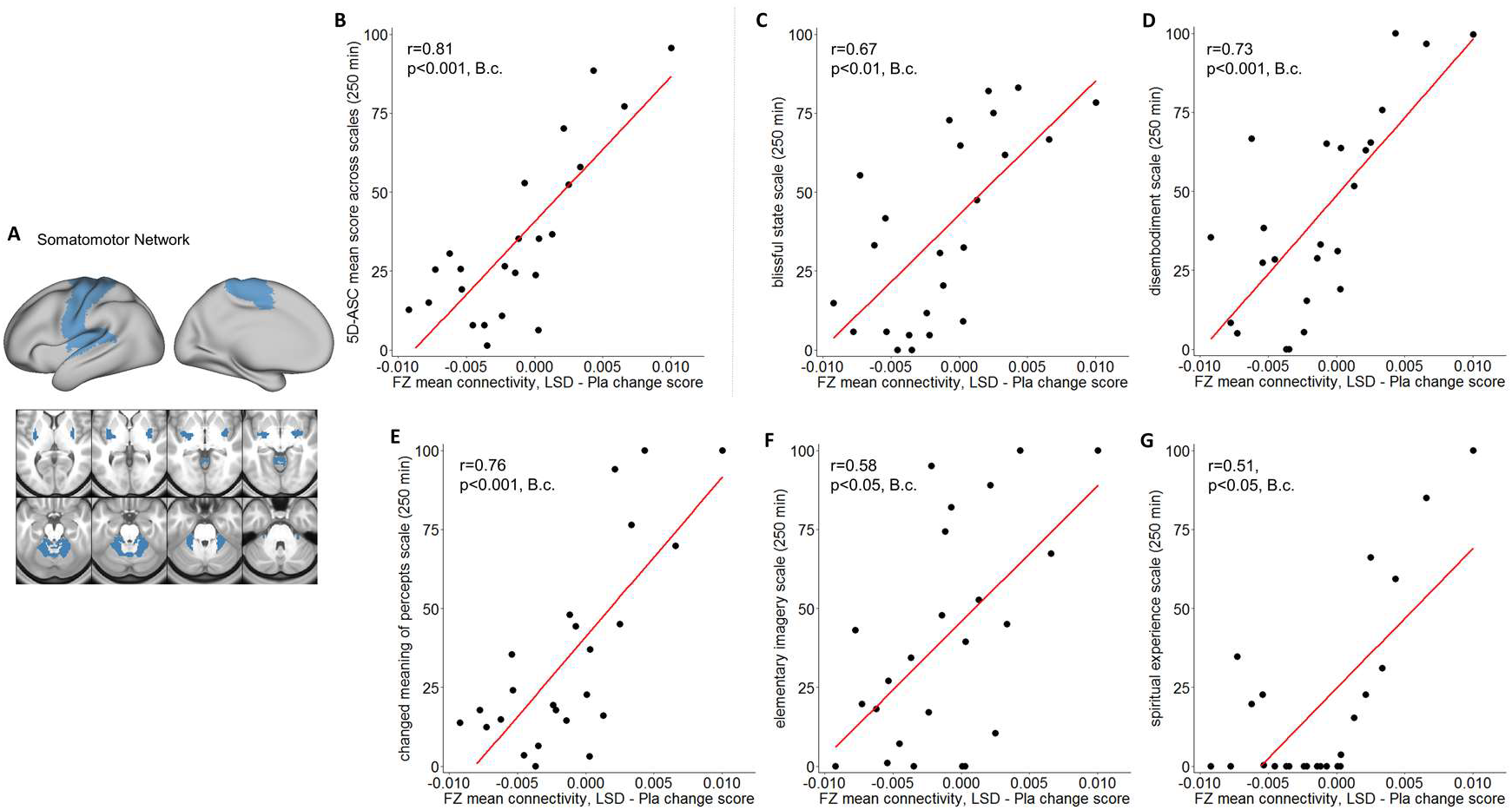
Correlation between Global Brain Connectivity and Subjective Effects. **(A)** The brain map illustrates lateral, medial, and subcortical view of the somatomotor network. **(B)** The scatterplot shows the significant positive correlation between Fz mean connectivity change (LSD – Pla condition, session 2, with GSR) in the somatomotor network and the mean 5-DASC short version score at 250 mins. **(C-G)** Scatterplots show the positive correlation between Fz mean connectivity change (LSD – Pla condition, session 2, with GSR) in the somatomotor network and the five subscales of the 5-DASC short version score at 250 mins: blissful state, disembodiment, changed meaning of percepts, elementary imagery, spiritual experience. B.c.: Bonferroni corrected. N=24.

Correlations between mean 5-DASC score and Fz connectivity in the other 6 networks and did not reveal significant relationships (all p>0.16, Bonferroni corrected). To further investigate the contribution of specific LSD-induced symptoms to the relationship with somatomotor network Fz connectivity, we calculated the correlation between Fz mean connectivity change in the somatomotor network with each 5-DASC short version scale separately. All five scale scores (blissful state, disembodiment, changed meaning of percepts, elementary imagery, spiritual experience) were significantly correlated with Fz mean connectivity change in the somatomotor network (all p<0.05, Bonferroni corrected, **Fig. 6 C-G**), indicating that the relationship between somatomotor network Fz connectivity and subjective effects was not driven by a specific LSD-induced symptom alone. Correlating mean Fz connectivity changes without GSR in the seven functional networks with subjective effects did not reveal any significant result (all p>0.3, unc.).

### GBC maps with GSR correlate predominantly with HTR2A and HTR7 cortical gene expression maps

LSD stimulates not only 5-HT2A R but also 5-HT-2C, -1A/B, -6, and -7 and dopamine D2 and D1 Rs. These receptors are differentially expressed across the cortex. To further investigate LSD’s receptor pharmacology, we tested the correlation between unthresholded Z-score map for LSD condition vs. Ketanserin+LSD condition + Placebo condition with and without GSR and six available receptor gene expression maps of interest (DRD1, DRD2, HTR1A, HTR2A, HTR2C, and HTR7) derived from the Allen Human Brain Atlas (29, 35). **Fig. 7A** shows the average GBC Z-score with and without GSR and the mean gene expression of the genes of interest within the seven functionally-defined networks using parcellations derived by Yeo et al. (32), Buckner et al. (33) and Choi et al. (34) (see also **Fig. 5**), indicating that gene expression is distinct by network. Next, we investigated whether there is a common pattern of distribution between the six gene expression maps. Correlation analyses showed that the expression of the main gene of interest (HTR2A) is highly negatively correlated with the expression of HTR7 (r=-0.68, p<0.001, Bonferoni corrected, **Fig. 7B**). **Fig. 7C** illustrates the cortical distribution of HTR2A gene expression. This HTR2A cortical gene expression map is highly correlated with the unthresholded GBC Z-score map for the LSD condition vs. Ketanserin+LSD condition + Placebo condition with GSR (r=0.50, p<0.001) , and higher than all other candidate serotonin receptor genes. While the correlation between the unthresholded GBC Z-score map without GSR and the HTR2A cortical gene expression map also reached significance (r=0.18, p<0.001), this correlation was significantly weaker than between the Z-score map with GSR and the HTR2A gene expression map (p<0.05, Bonferroni corrected, **Fig. 7F**). Taking into account all available gene expression maps the correlation between the Z-score map with GSR and the HTR2A gene expression map was higher that 95.9% of all possible correlations. The GBC Z-score map with GSR and the HTR7 gene expression map was lower than 99.8% of all possible correlations, indicating a strong negative relationship (r=-0.63, p<0.001, **Fig. 7D**). The correlation between both HTR2A and HTR7 with the GBC Z-Score map is not surprising considering the strong negative correlation between HTR2A and HTR7 gene expression maps (**Fig 7B**). Lastly, **Fig 7F** illustrates that correlation coefficients between gene expression maps and GBC Z-score maps were significantly stronger with GSR (all p<0.05, Bonferoni corrected), except for DRD1 expression map, where the absolute value of the correlation coefficient increased when correlated with the Z-score map without GSR.

**Fig. 7.**
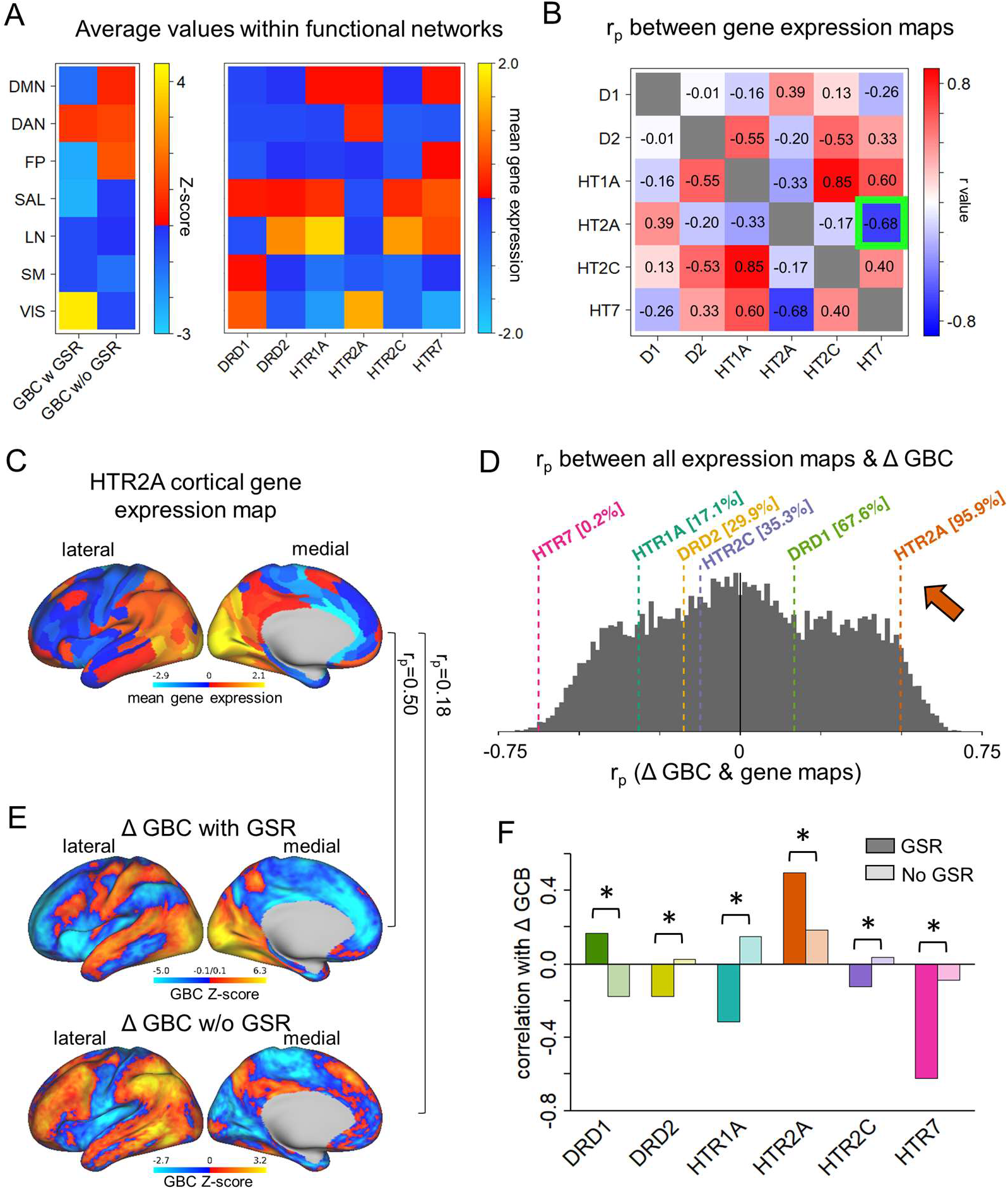
Correlation between GBC and cortical gene expression maps. **(A)** The top left panel shows the average GBC Z-score (LSD condition vs. Ketanserin+LSD condition + Placebo condition) with and without GSR and mean gene expression value within functional networks. **(B)** The top right panel shows the correlation (Pearson’s r) between the gene expression maps, highlighting (green) the negative correlation between the expression maps of HTR2A and HTR7. **(C)** The brain map illustrates the cortical expression levels (Z-score) of HTR2A in the left hemisphere (lateral and medial view). **(D)** The histogram depicts the correlation between all gene expression maps and the unthresholded Z-score map for LSD condition vs. Ketanserin+LSD condition + Placebo condition) with GSR. The colored lines highlight the gene expression maps of interest. **(E)** Unthresholded Z-score map for LSD condition vs. Ketanserin+LSD condition + Placebo with (top) and without (bottom) GSR. Red/orange areas indicate regions where participants exhibited stronger GBC in the LSD condition, whereas blue areas indicate regions where participants exhibited reduced GBC in the LSD condition, compared with (Ketanserin+LSD) + Placebo. r_p_ values are the respective correlation coefficients between Z-score maps and HTR2A gene expression map. **(F)** The bar graph shows the correlation coefficients (Pearson’s r) between each gene expression map of interest and Z-score maps for LSD condition vs. Ketanserin+LSD condition + Placebo condition with and without GSR. * indicates significant difference between correlations between Z-score map with and without GSR and gene expression map, p<0.05, Bonferroni corrected.

## DISCUSSION

Interest in the potential clinical effects of psychedelics is boosted by positive preliminary reports on the safety and tolerability in healthy participants as well as patient populations (5, 6, 36, 37). However, the underlying pharmacology and neurobiology of psychedelics, in particular the prototypical LSD, is scarcely investigated in humans. The current study closes major knowledge gaps by showing that i) LSD increases GBC across sensory functional networks and reduces GBC in associative networks, which is sensitive to GS removal; ii) time-dependent effects are only found in the interaction with katenserin; iii) GBC in the somatomotor network predicted subjective effects; iv) LSD-induced effects on GBC and subjective symptoms are linked to the pharmacology of the 5-HT2A receptor; v) innovative gene expression analyses across cortex reveal for the first time a correspondence between specific spatial expression patterns and in-vivo pharmacological effects in humans.

### LSD increases GBC across sensory functional networks and reduces GBC in associative networks

Comparing LSD to (Ket+LSD)+Pla conditions shows that LSD induces hyper-connectivity predominantly in sensory areas, i.e. the occipital cortex, the superior temporal gyrus, and the postcentral gyrus, as well as the precuneus. Hypo-connectivity was induced in subcortical areas as well as cortical areas associated with associative networks, such the medial and lateral prefrontal cortex, the cingulum, the insula, and the temporoparietal junction (**Fig. 1**). These results indicate that LSD causes a de-synchronization of associative networks whereas sensory areas are more highly integrated (18). This is at least partially in line with previous studies reporting increased seed-based V1 resting-state connectivity with the rest of the brain after LSD administration, (17) decreases of connectivity within the DMN after ayahuasca intake (38), and desynchronization within the DMN after psilocybin infusion measured with magnetoencephalography (39). Subcortical areas predominantly show hypoconnectivity. However, the amygdala shows hyper-connectivity under LSD. Amygdala neurons abundantly express 5-HT2A receptors, and alterations in amygdala activity and connectivity have been hypothesized to be important for potential beneficial clinical effects of psychedelics (40–42). Furthermore, our analysis showed that participants with the highest LSD-induced coupling within sensory networks also showed the strongest LSD-induced de-coupling in associative networks (**Fig 2**). This suggests that LSD-induced alterations in information flow across these networks probably results from linked systems-level perturbations, as opposed to being due to separate and dissociable mechanisms across subjects. This pattern of hyper-and hypo-connectivity may therefore underlie the psychedelic state, suggesting increased processing of sensory information which is not counterbalanced by associative network integrity, therefore resulting in an altered state of consciousness whereby sensory experiences cannot be integrated, leading to psychedelic symptoms.

A virtually identical pattern of hyper-connectivity in sensory networks and hypo-connectivity in associative networks is revealed when contrasting LSD effects with the condition where LSD is blocked by katensirin. This result was virtually indistinguishable from LSD vs. Placebo condition. Since ketanserin has high antagonistic properties particularly on the 5-HT2A receptor, these results indicate that LSD-induced alterations in GBC are presumably highly dependent on stimulation of the 5-HT2A receptor (28). This is also supported by the results that pre-treatment of LSD with ketanserin induced only negligible changes compared to Placebo (**Fig S3**), indicating that ketanserin blocked virtually all LSD-induced alterations in GBC. This is in line with data on subjective LSD-induced effects which were normalized by the pretreatment with ketanserin (**Fig. 1F**). Therefore, our results extend previous studies by revealing a pattern of brain connectivity characterized by the integration of sensory networks and dis-integration of associative networks which is presumably underlying the LSD-induced altered state of consciousness. Furthermore, our data are pointing to the importance of the 5-HT2A receptor system in LSD-induced alterations, not only in subjective effects, but also in resting state functional connectivity.

### Time-dependent effects of LSD

Animal studies suggest distinct temporal phases of LSD pharmacology (30, 31). Therefore, we investigated the time-dependent effects of LSD on subjective effects as well as on GBC. As shown in **Fig. 3**, subjective effects were highest 180 min after LSD administration and decreased in intensity 250 and 360 min after administration as expected (26, 43). No differences were found between Placebo and Ket+LSD conditions at any time point, with subjective effects in both conditions being very low in intensity (<4.7%). This shows that ketanserin blocked subjective LSD-induced effects over the whole time course, indicating that subjective effects are most likely attributable to 5-HT2A receptor stimulation. To investigate the time course of LSD-induced effects on GBC, two resting-state scans were analyzed, conducted 75 minutes (session 1) and 300 minutes (session 2) after the second drug administration. While no significant differences were observed when comparing session 1 and 2 within the placebo and the LSD condition (**Fig. S4**), participants showed significant changes in GBC in session 2 compared to session 1. Taken together with time-dependent results observed in the functionally defined networks (**Fig. 5**), the blocking effect of ketanersin is particularly evident in the session 1 across all networks. Specifically, ketanserin not only blocks LSD effects in session 1 but also augments the effects seen in placebo, indicating the opposite mechanism of action from that seen by LSD – namely 5-HT2A receptor antagonism (28, 44). On the other hand, it seems possible that there exist two distinct pharmacolological time phases as described in animal studies. The first phase may be modulated by 5-HT2A receptor activation and the second phase possibly by D2 receptor activation, as suggested by preclinical work (30, 31). This hypothesis of time-dependent complex receptor pharmacology awaits further testing. First, studies are needed to investigate the effect of ketanserin alone on GBC to verify the preferential effects of 5-HT2A antagonism. Second, studies using pre-treatment of LSD by antagonists on receptors other than 5-HT2A are needed to determine if the second phase is indeed modulated by another receptor system. Lastly, indications of different pharmacological phases are not evident from subjective drug effects which remain completely blocked by ketanserin. Studies using higher doses of LSD are therefore needed to investigate if the potential effect of LSD’s action on other receptors becomes more pronounced and therefore subjectively accessible.

### LSD’s effect on GBC is sensitive to GS removal

One small study by Tagliazucchi et al. has previously investigated the effects of intravenously administered LSD on GBC and reports that highlevel association cortices (partially overlapping with the default-mode, salience, and frontoparietal attention networks) and the thalamus showed increased GBC under LSD (20). These results are partially contradictory to the results presented in **Fig. 1.** However, this previous study did not take into account the influence of the global signal (GS) which can contain large amounts of non-neuronal artifact (e.g. respiration, (21, 45)) and induce spuriously high correlations across the brain (11, 25). Due to these methodological discrepancies, we re-analyzed our data without applying GS regression in concordance with Tagliazucchi et al. (20). This analysis revealed connectivity patterns at least partially overlapping with the results by Tagliazucchi et al. (20), i.e. increased GBC in fronto-parietal, temporal, and subcortical areas (**Fig. S5**). However, without GSR, no significant differences between LSD and Placebo were found in the seven functionally defined networks (**Fig. 5**). In stark contrast to the analysis that removed global artifact, the analysis without GSR showed a positive correlation between hyper- and hypo-connectivity change scores, indicating that participants with the highest LSD-induced hyperconnectivity showed the weakest LSD-induced de-coupling. Furthermore, connectivity values with GSR and those without GSR were not significantly correlated within subjects (**Fig. S6**). This suggests that differing results between analyses that removed artifact and those that did not cannot be explained by a mean-shift in connectivity values, but are due to an independent process within each subject. A likely explanation is that GSR removes non-neural artifacts and therefore provides a method to better isolate functional networks in pharmacological resting-state connectivity studies (11, 25). This interpretation is strongly supported by the findings that subjective effects did not correlate with Fz connectivity changes without GSR. Finally, spatial correlations with gene expression maps (discussed below) were significantly attenuated for results that did not remove GS.

### LSD-induced alterations in GBC in the somatomotor network are associated with subjective effects

The LSD-induced change in connectivity in the somatomotor network correlated significantly with general and specific subjective LSD-induced effects (mean across all scales, blissful state, disembodiment, changed meaning of percepts, elementary imagery, spiritual experience). Participants with increased connectivity in the somatomotor network also showed higher subjective effects. On average, the change in connectivity between the LSD and Placebo condition in the somatomotor network was not significant. However, this is likely explained by the heterogeneous connectivity changes within this network: while the pre- and postcentral gyrus predominantly showed increases in GBC, medial areas were hypo-connected. Connectivity changes in other functional networks were not significantly correlated with subjective effects. This points to the importance somatomotor network brain regions and their connectivity with the rest of the brain for psychedelic experiences. This is in line with previous results obtained from task-related data showing that the supplementary motor area is associated with LSD-induced alterations in meaning and personal relevance processing (27). These results also support broader theories of consciousness emphasizing the importance of the sensorimotor system for the perception of presence and agency, and therefore a sense of self (e.g., interoceptive predictive coding model of conscious presence (46), comparator model (47)) (48, 49). Furthermore, alterations in sensorimotor gating have been suggested to underlie psychedelic experiences (50–52). Somatomotor system activity and connectivity has also been implicated in the pathophysiology of schizophrenia (53), an illness characterized by delusions and alterations in the sense of self, potentially arising from alterations in sensorimotor gating deficits in an inferential mechanism that allows distinguishing whether or not a sensory event has been self-produced (54). The current results corroborate and extend these previous findings by showing that somatomotor network connectivity is also closely associated with an LSD-induced psychedelic state.

### LSD-induced alterations in GBC correlate with HTR2A and HTR7 cortical gene expression

To further investigate LSD’s receptor pharmacology we specifically used the threshold-free Z-score map of LSD effects relative to ketanserin blockade. The logic here is that such a map may reflect ketanserin-specific contributions to LSD blockade, which is hypothesized to involve the HTR2A receptor. This map was then correlated with gene expression maps of receptors that may be stimulated by LSD (3). LSD-induced changes in functional connectivity after GSR exhibited strong positive relationships with HTR2A expression (higher than 95.9% of all possible gene expression correlations, **Fig. 7**). These results show that LSD-induced changes in GBC quantitatively match the spatial expression profile of genes coding for the 5-HT2A receptor, supporting the central role of this receptor system in LSD’s neuronal and subjective effects. LSD-induced changes in functional connectivity were also highly negatively correlated with HTR7 gene expression (lower than 99.8% of all possible gene expression correlations, **Fig. 7**). This can be explained by the highly anti-correlated expression of these two genes (**Fig. 7**). However, it is also possible that the 5-HT7 receptor functionally contributes to LSD-induced effects. In contrast to its agonistic properties on the 5-HT2A receptor, LSD has been reported to be an antagonist in the 5-HT7 receptor (55). Since previous studies have shown that 5-HT7 receptor antagonists have anti-psychotic potential (56–59), it seems very unlikely that LSD’s effects have a strong and appreciable contribution on the 5-HT7 receptor. However, future studies should examine 5-HT7 receptor pharmacology more carefully as they may reveal a role of this receptor system in procognitive effects that contrast those of LSD. Finally, we show that the spatial match between gene expression maps and GBC maps is significantly improved after artifact removal, pinpointing the important role of careful BOLD signal de-noising in pharmacological imaging. These results also highlight the validity of this approach as a general method for relating spatial gene expression profiles to neuropharmacological manipulations in humans.

### Conclusion

In summary, the current results close major knowledge-gaps regarding the neurobiology and neuropharmacology of LSD. First, we show that LSD induces widespread alterations of GBC in cortical and subcortical brain areas, characterized by a synchronization of sensory functional networks and dis-integration of associative networks. We show that this effect is sensitive to artifact removal, which has important implications for future pharmacological rsfMRI studies. Second, we investigated the receptor-pharmacology of LSD, showing that the 5-HT2A receptor plays a critical role in subjective and neuronal LSD-induced effects. However, analyzing the time course of LSD-induced alterations in functional connectivity, it seems likely that at a later phase, modulation by receptors other than the 5-HT2A receptor is involved. The comparison of LSD-induced effects on functional connectivity and receptor-gene expression maps underscores the interpretations of 5-HT2A pharmacology and points to potentially impactful studies on 5-HT7 receptor pharmacology. Lastly, in line with various theories of consciousness we showed that the somatomotor system in particular is related to LSD-induced psychedelic effects. Collectively, these results deepen our understanding of psychedelic compounds and offer important directions for development of novel therapeutics.

## METHODS

### Participants

See supplemental methods.

### Study Design

The study employed a fully double-blind, randomized, cross-over design (see **Fig. S1**). Specifically, participants received either:

i. placebo+placebo (Pla) condition: placebo (179mg Mannitol and Aerosil 1mg po) after pretreatment with placebo (179mg Mannitol and Aerosil 1mg po);
ii. placebo + LSD (LSD) condition: LSD (100 μg po) after pretreatment with placebo (179mg Mannitol and Aerosil 1mg po), or
iii. ketanserin+LSD (Ket+LSD) condition: LSD (100 μg po) after pretreatment with the 5-HT2A antagonist ketanserin (40 mg po) at three different occasions two weeks apart.

Pretreatment with placebo or ketanserin occurred 60 min before treatment with placebo or LSD. The resting-state scan was conducted 75 and 300 min after treatment administration. Participants were asked to not engage in repetitive thoughts such as counting and close their eyes during the resting state scan. Compliance to this instruction was monitored online using eye tracking (NordicNeuroLab VisualSystem, http://www.nordicneurolab.com/). The 5D-ASC (a retrospective self-report questionnaire) (60) was administered to participants 720 min after drug treatment to assess subjective experience after drug intake. In addition, a short version of the 5D-ASC was administered 180, 250, and 360 min after drug treatment to assess the time course of subjective experience. Participants were required to abstain from smoking for at least 60 min before MRI assessment and from drinking caffeine during the test day.

### Neuroimaging Data Acquisition, Preprocessing

See supplemental methods.

### Global Brain Connectivity Calculation

Most connectivity studies focus on pre-defined areas (i.e. seed-based approaches). Such approaches assume ‘dysconnectivity’ across similar regions or networks. However, functional dysconnectivity induced by LSD, especially across heterogeneous associative cortical circuits, may exhibit variability across people. To address this, here we applied recently optimized neuroimaging analytic techniques to identify dysconnectivity in a data-driven fashion, termed global brain connectivity (GBC)(61–63). GBC is a measure that examines connectivity from a given voxel (or area) to all other voxels (or areas) simultaneously by computing average connectivity strength – thereby producing an unbiased approach as to the location of dysconnectivity. Also, unlike typical seed approaches, GBC involves one statistical test per voxel (or area) rather than one test per voxel-to-voxel pairing, substantially reducing multiple comparisons (e.g., 30,000 rather than ∽450 million tests). These improvements dramatically increase the chances of identifying pharmacologically-induced dysconnectivity, or individual differences correlated with symptoms, as we demonstrated by our prior studies conducted in clinical populations (19, 61, 63, 64). By extension, this approach can be readily applied to pharmacological neuroimaging studies. Specifically, the GBC approach (19, 63) was applied using in-house Matlab tools studies (61–63), extended across all gray matter voxels in the brain, as defined via the CIFTI image space, which was obtained via an adapted version of FreeSurfer software fine-tuned by the HCP pipelines (65). Finally, for each grayordinate in the CIFTI image space, we computed a correlation with every other whole-brain grayordinate, transformed the correlations to Fisher z-values, and finally computed their mean. This calculation yielded a GBC map for each subject where each voxel value represents the mean connectivity of that voxel with all other gray matter voxels in the brain. We also verified that differences in variance of BOLD signals did not drive our GBC results, as predicted by our prior computational modeling work (66). To this end, we computed GBC using a non-normalized covariance measure, which did not alter effects. Appropriate whole-brain type I error correction was computed via FSL’s PALM tool (see **2^nd^-Level Group Comparisons** below).

### Global Signal Regression

There still remains considerable controversy regarding the utility of mean signal de-noising strategies (21, 22), with clear pros/cons. GSR was performed using standard procedures by calculating mean raw BOLD signal averaged over all gray matter voxels for each time point, explicitly excluding ventricles and white matter (which are defined as separate nuisance regressors). The GS and its first derivative (with respect to time) were used as nuisance predictor terms within a multiple linear regression model along with other nuisance predictor terms (ventricular signal, white matter signal, movement parameters, and the first derivatives of each of these, as noted above). Because of emerging findings suggesting that clinical populations exhibit elevated GS variability (66), we separately examined results without GSR implemented. This demonstration is particularly important given recent reports suggesting that the GS may be abnormally altered in specific clinical populations (66, 67), but also that it may contain major elements of respiratory artifact (21), which could influence GBC analyses.

### Quality Assurance Analyses, 2^nd^ Level Statistical Analysis, Statistical Analysis of Behavioral Data

See supplemental methods and **Fig. S2.**

### Gene expression preprocessing

To relate LSD-related neuroimaging effects to the cortical topography of gene expression for candidate receptors, we used the Allen Human Brain Atlas (AHBA). The AHBA is a publicly available transcriptional atlas containing gene expression data, measured with DNA microarrays, that were sampled from hundreds of histologically validated neuroanatomical structures across six normal post-mortem human brains (35). All reported analyses were performed on group-averaged gene expression maps in the left cortical hemisphere, which were generated following a previously reported procedure (29). In brief, a group-averaged, dense cortical expression map was constructed through a neurobiologically informed approach using a surface-based Voronoi tessellation combined with a 180-area unilateral parcellation with the Human Connectome Project’s Multi-Modal Parcellation (MMP1.0) (68).

## ACKNOWLEDGMENTS

This research was financially supported by grants from the Swiss National Science Foundation (SNSF, P2ZHP1_161626), the Swiss Neuromatrix Foundation (2015-0103), and the Usona Institute (2015-2056).

## SUPPLEMENTAL INFORMATION

### Supplemental Methods

#### Participants

Participants were recruited through advertisements placed in local universities and underwent a screening visit before inclusion in the larger study protocol (1). All included subjects were healthy according to medical history, physical examination, blood analysis, and electrocardiography. The Mini-International Neuropsychiatric Interview (MINI-SCID) (2), the DSM-IV self-rating questionnaire for Axis-II personality disorders (SCID-II) (3), and the Hopkins Symptom Checklist (SCL-90-R) (4) were used to exclude subjects with present or previous psychiatric disorders or a history of major psychiatric disorders in first-degree relatives. Participants were asked to abstain from the use of any prescription or illicit drugs for a minimum of two weeks prior to the first test day and for the duration of the entire study, and to abstain from drinking alcohol for at least 24 h prior to test days. Urine tests and self-report questionnaires were used to verify the absence of drug and alcohol use. Urine tests were also used to exclude pregnancy. Further exclusion criteria included left-handedness, poor knowledge of the German language, cardiovascular disease, history of head injury or neurological disorder, history of alcohol or illicit drug dependence, magnetic resonance imaging (MRI) exclusion criteria including claustrophobia, and previous significant adverse reactions to a hallucinogenic drug.

Twenty-five participants took part in the study. One subject was excluded due to poor neuroimaging signal quality. Therefore a sample of 24 participants was included in the final analysis (n=19 males and n=5 females; mean age = 25.00 years; standard deviation (SD) = 3.60 years; range 20–34 years). All participants provided written informed consent statements in accordance with the declaration of Helsinki before participation in the study. Subjects received written and oral descriptions of the study procedures, as well as details regarding the effects and possible risks of LSD and ketanserin treatment. The Swiss Federal Office of Public Health, Bern, Switzerland, authorized the use of LSD in humans, and the study was approved by the Cantonal Ethics Committee of Zurich. The study was registered at ClinicalTrials.gov (NCT02451072).

#### Neuroimaging Data Acquisition

Magnetic resonance imaging (MRI) data were acquired on a Philips Achieva 3.0T whole-body scanner (Best, The Netherlands). A 32-channel receive head coil and MultiTransmit parallel radio frequency transmission was used. Images were acquired using a whole-brain gradient-echo planar imaging (EPI) sequence (repetition time= 2,500ms; echo time=27 ms; slice thickness=3mm; 45 axial slices; no slice gap; field of view = 240×240 mm^2^; in-plane resolution = 3×3 mm; sensitivity-encoding reduction factor = 2.0). Additionally, two high-resolution anatomical images (voxel size = 0.7mm^3^) were acquired using T1-weighted and T2-weighted sequences.

#### Preprocessing

Structural and functional MRI data were first preprocessed according the methods provided by the Human Connectome Project (HCP), outlined below, and described in detail by the WU-Minn HCP consortium (5). These open-source HCP algorithms, optimized for our specific acquisition parameters and Yale’s High Performance Computing resources, represent the current state-of-the-art in distortion correction, registration, and maximization of high-resolution signal-to-noise (SNR). Here we briefly describe the processing steps. Complete details are outlined by Glasser and colleagues (5). First, the T1w/T2w images were corrected for bias-field distortions and warped to the standard Montreal Neurological Institute-152 (MNI-152) brain template through a combination of linear and non-linear transformations using the FMRIB Software Library (FSL) linear image registration tool (FLIRT) and non-linear image registration tool (FNIRT) (6). Then, FreeSurfer’s recon-all pipeline was employed to compute brain-extraction, within-subjects registration, and individual cortical and subcortical anatomical segmentation (7). T1w/T2w images were then converted to the Connectivity Informatics Technology Initiative (CIFTI) volume/surface ‘grayordinate’ space.

Raw BOLD images were first corrected for field inhomogeneity distortion, phase encoding direction distortions and susceptibility artifacts using the pair of reverse phase-encoded spin-echo field-map images implemented via FSL’s TOPUP algorithm (8). Motion-correction was then performed by registering each volume in a run to the corresponding single-band reference image collected at the start of each run. BOLD images were then registered to the structural images via FLIRT/FNIRT, and a brain-mask was applied to exclude signal from non-brain tissue. After processing in NIFTI volume space, BOLD data were converted to the CIFTI gray matter matrix by sampling from the anatomically-defined gray matter ribbon.

Following these minimal HCP preprocessing steps, a high-pass filter (>0.5 Hz) was applied to the BOLD time series in order to remove low frequencies and scanner drift. In-house MATLAB tools were used to compute the average variation in BOLD signal in the ventricles and deep white matter. This signal was regressed out of the gray matter time series as a nuisance variable because any BOLD signal change in those structures was likely due to pervasive rather than cortical activity. Finally, mean gray matter time series (i.e. the global signal) was also regressed to address spatially pervasive artifact, such as respiration. Finally, all data were motion-scrubbed as recommended by Power et al. (9, 10). As accomplished previously (11), all image frames with possible movement-induced artifactual fluctuations in intensity were identified via two criteria: first, frames in which the sum of the displacement across all 6 rigid body movement correction parameters exceeded 0.5mm (assuming 50mm cortical sphere radius) were identified. Second, root mean square (RMS) of differences in intensity between the current and preceding frame was computed across all voxels and divided by mean intensity. Frames in which normalized RMS exceeded 1.6 times the median across scans were identified. The frames flagged by either criterion, as well as the one frame preceding and two frames following each flagged frame, were marked for exclusion (logical or). Subjects with more than 50% frames flagged were completely excluded from all analyses. All the included subjects in the final samples passed these criteria.

#### Quality Assurance Analyses

For quality assurance purposes we computed the following measures: i) signal-to-noise ratio (defined as mean signal over the entire BOLD time series for a given voxel divided by its standard deviation), and ii) the percentage of ‘scrubbed’ images. In turn, we correlated these measures with mean Fz-connectivity with and without GSR for the first and second session in the LSD condition (**Fig. S2**). All correlations were non-significant indicating that changes in GBC induced by LSD are not attributable to motion and image artifacts.

#### 2^nd^ Level Statistical Analysis

GBC maps for each subject, condition, and session were entered into a 2x3 repeated-measures ANOVA and tested using FSL’s Permutation Analysis of Linear Models (PALM, (12)). Threshold-free cluster enhancement (TFCE) was used to avoid the need to define clusters using arbitrary thresholds for cluster size. The statistical images were thresholded at p<0.05 (family-wise error corrected), with 10000 permutations. For further analysis connectivity strength (Fz) values for hyper- and hypo-connected areas (based on the LSD vs (Ketanserin+LSD)+Placebo contrast) were averaged across voxels for each participant and condition. Furthermore, Fz values for voxels within seven functionally-defined networks using parcellations derived by Yeo et al. (13), Buckner et al. (14) and Choi et al. (15) were averaged for each participant and condition. All analyses were performed with and without GSR. Results were visualized using the Connectome Workbench software (https://www.humanconnectome.org/software/connectome-workbench.html).

#### Statistical Analysis of Behavioral Data

The 5D-ASC comprises 94 items that are answered on visual analogue scales (16). Scores were calculated for 11 recently validated scales (17): experience of unity, spiritual experience, blissful state, insightfulness, disembodiment, impaired control and cognition, anxiety, complex imagery, elementary imagery, audio-visual synesthesia, and changed meaning of percepts. The short version of the 5D-ASC includes the 45 items that comprise the spiritual experience, blissful state, disembodiment, elementary imagery, and changed meaning of percepts scales. 5D-ASC score was analyzed using a repeated-measures ANOVA with treatment condition (Pla, LSD, and Ket+LSD) and scale as within-subject factors. 5D-ASC short-version score was analyzed using a repeated-measures ANOVA with treatment condition (Pla, LSD, and Ket+LSD), scale, and time (180, 250, and 360 min) as within-subject factors. The 5D-ASC short-version scores of one participant could not be analyzed due to missing data at 360 min after administration. Bonferroni-corrected Pearson correlations were conducted to investigate the relationship between Fz values within the seven functionally defined networks at session 2 and subjective drug effects (5D-ASC short version at 250 min).

**Figure S1.**
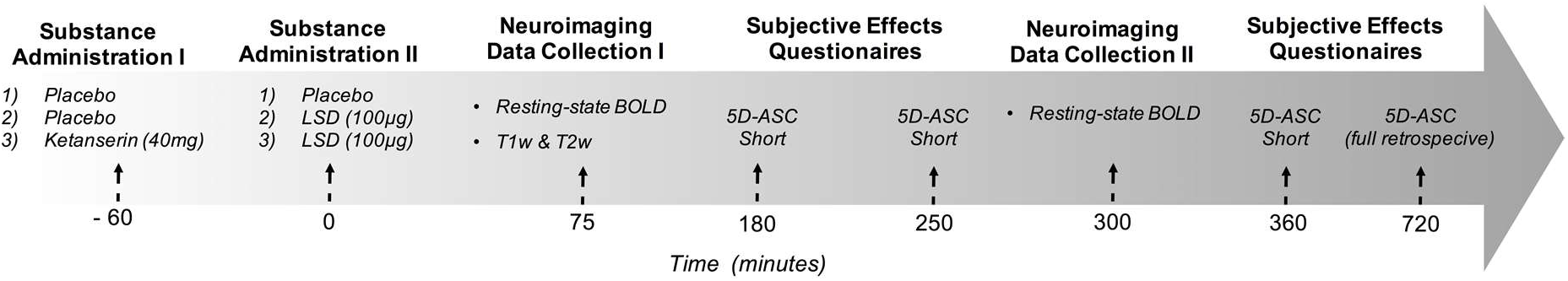
Study Design. The study employed a fully double-blind, randomized, within-subject cross-over design. 25 total participants received either: 1) placebo + placebo (Pla) condition: placebo (179mg Mannitol and Aerosil 1mg po) after pretreatment with placebo (179mg Mannitol and Aerosil 1mg po); 2) placebo + LSD (LSD) condition: LSD (100 μg po) after pretreatment with placebo (179mg Mannitol and Aerosil 1mg po), or 3) ketanserin+LSD (Ket+LSD) condition: LSD (100 μg po) after pretreatment with the 5-HT2A antagonist ketanserin (40 mg po) in a randomized balanced order at three different occasions two weeks apart.

**Figure S2.**
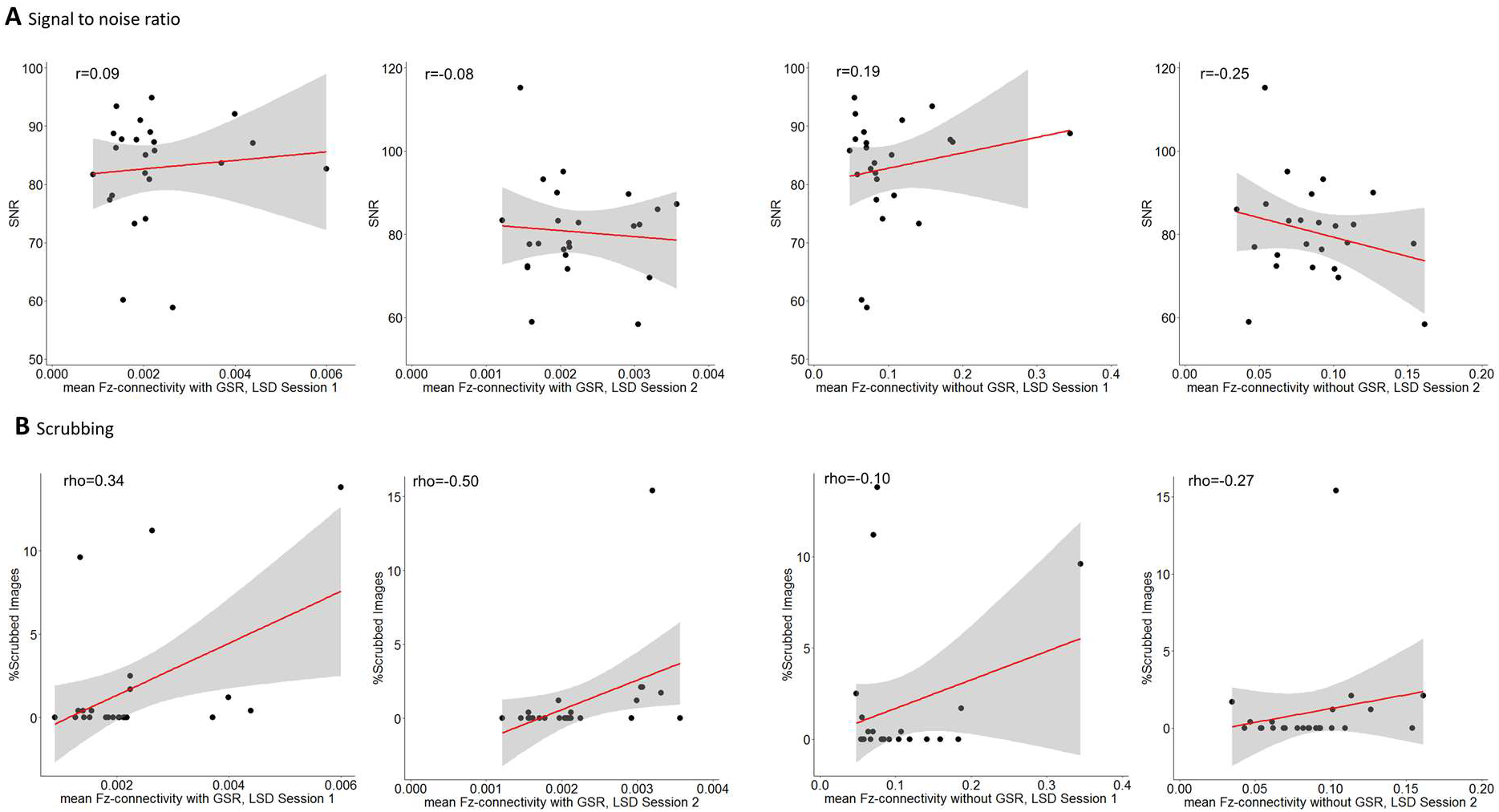
Quality Control (QC) Measures Do Not Correlate with Mean GBC. We computed the relationship between QC measures and the mean GBC for various conditions across subjects. **(A)** Signal-to-noise ratio (defined as mean signal over the entire BOLD time series for a given voxel divided by its standard deviation). **(B)** The percentage of ‘scrubbed’ images. Here we used the following criterion to compute a percentage of frames flagged for high head motion: First, frames in which sum of the displacement across all 6 rigid-body movement correction parameters >0.5 mm (assuming 50 mm cortical sphere radius) were identified. Secondly, root mean square (RMS) of differences in intensity between the current and preceding frame was computed across all voxels and divided by mean intensity. Frames in which normalized RMS exceeded the value of 3 were identified. The frames flagged by either criterion were marked for exclusion, as well as the one preceding and 2 frames following the flagged frame. Subjects with >50% frames flagged were completely excluded from analyses. We quantified these QC measures in relation to mean Fz-connectivity with and without GSR for the first and second session in the LSD condition. None of the correlations were significant indicating that changes in GBC induced by LSD are not attributable to motion and image artifacts.

**Figure S3.**
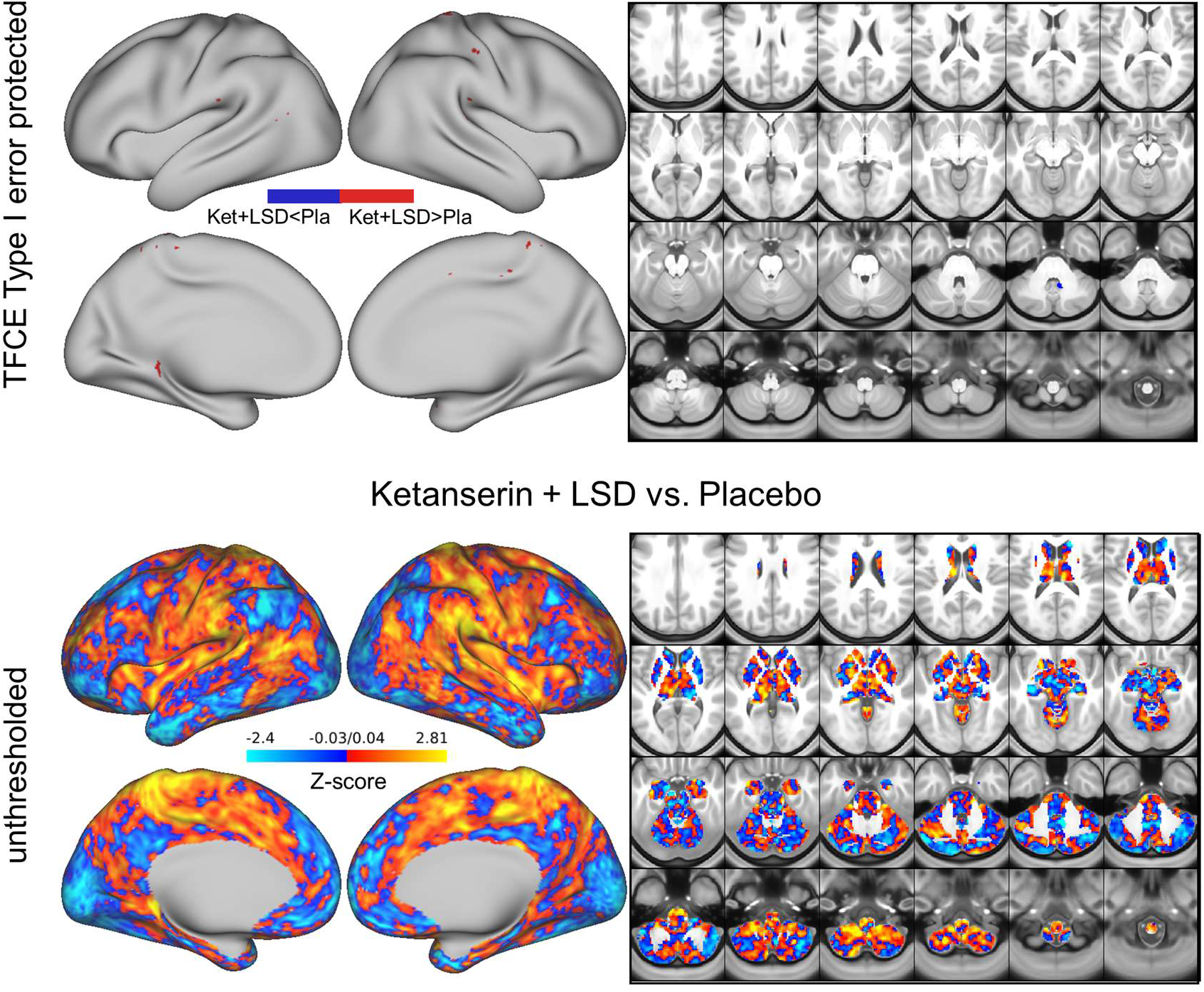
Effect of Ketanserin+LSD vs. Placebo on GBC. Top panel displays significant (TFCE type 1 error protected) areas showing increased (red) and decreased (blue) GBC in the Ket+LSD condition compared to Placebo, which were trivial. Lower panel shows the corresponding unthresholded Z-score map. Red/orange areas indicate regions where participants exhibited stronger GBC in the Ket+LSD condition, whereas blue areas indicate regions where participants exhibited reduced GBC in the Ket+LSD condition, compared with Placebo condition.

**Figure S4.**
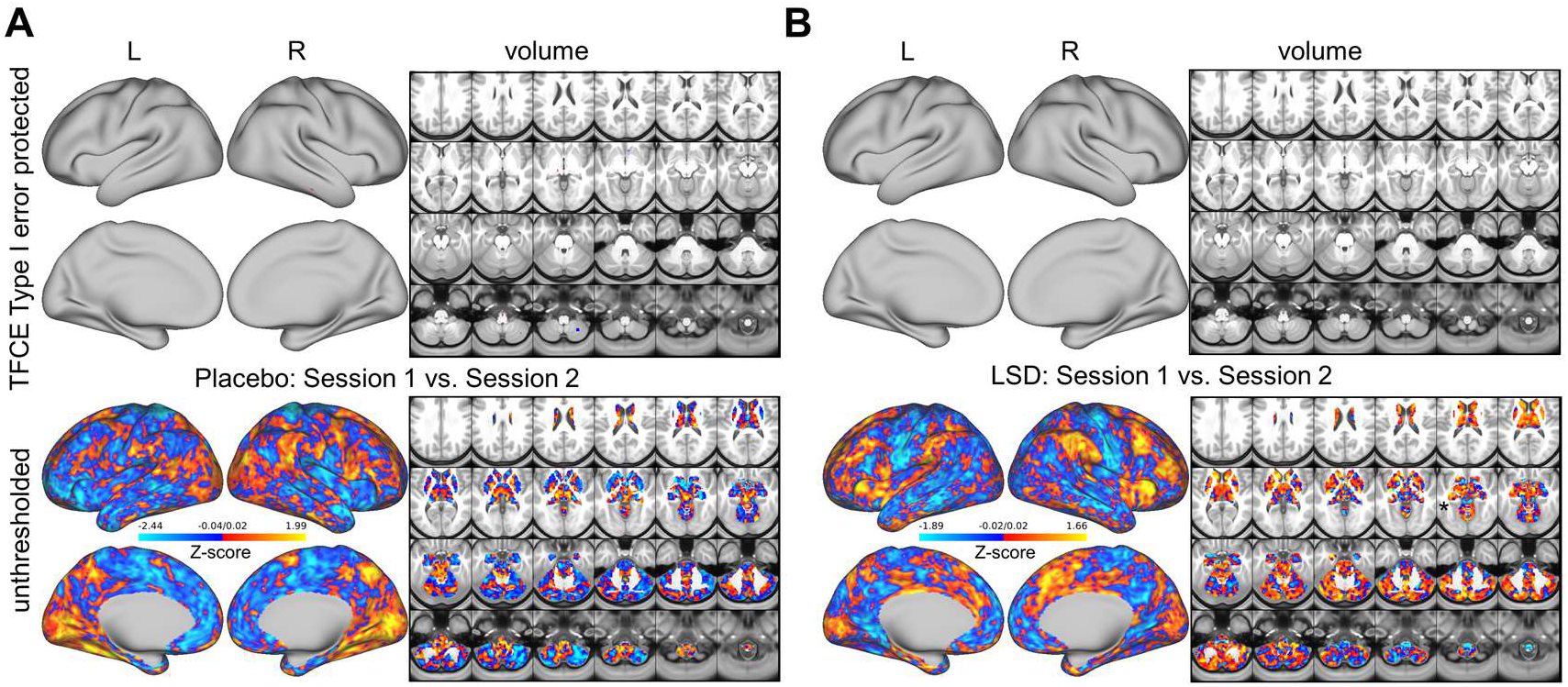
Effect of Session on Global Brain Connectivity. Animal studies suggest distinct temporal phases of LSD pharmacology, with an early phase mediated by 5-HT2A and a later phase mediated by D2 receptor stimulation (18, 19). However, temporal differences in receptor contributions to the effects of LSD have not been investigated in humans. Therefore, the current study quantified GBC, with two independent scans, at two distinct time points (75 and 300 min after administration) in a sample of 24 participants who each underwent three drug treatment conditions: 1) placebo after pretreatment with placebo (Pla), 2) LSD after pretreatment with placebo (LSD), and 3) LSD after pretreatment with ketanserin (Ket+LSD). **(A)** Placebo condition. **(B)** LSD condition. Top panel displays no significant (TFCE type 1 error protected) areas showing increased or decreased GBC in session 1 (75 minutes after second drug administration) compared to session 2 (300 minutes after second drug administration). Lower panel shows the corresponding unthresholded Z-score map. Red/orange areas indicate regions where participants exhibited stronger GBC in session 1, whereas blue areas indicate regions where participants exhibited reduced GBC in session 2.

**Figure S5.**
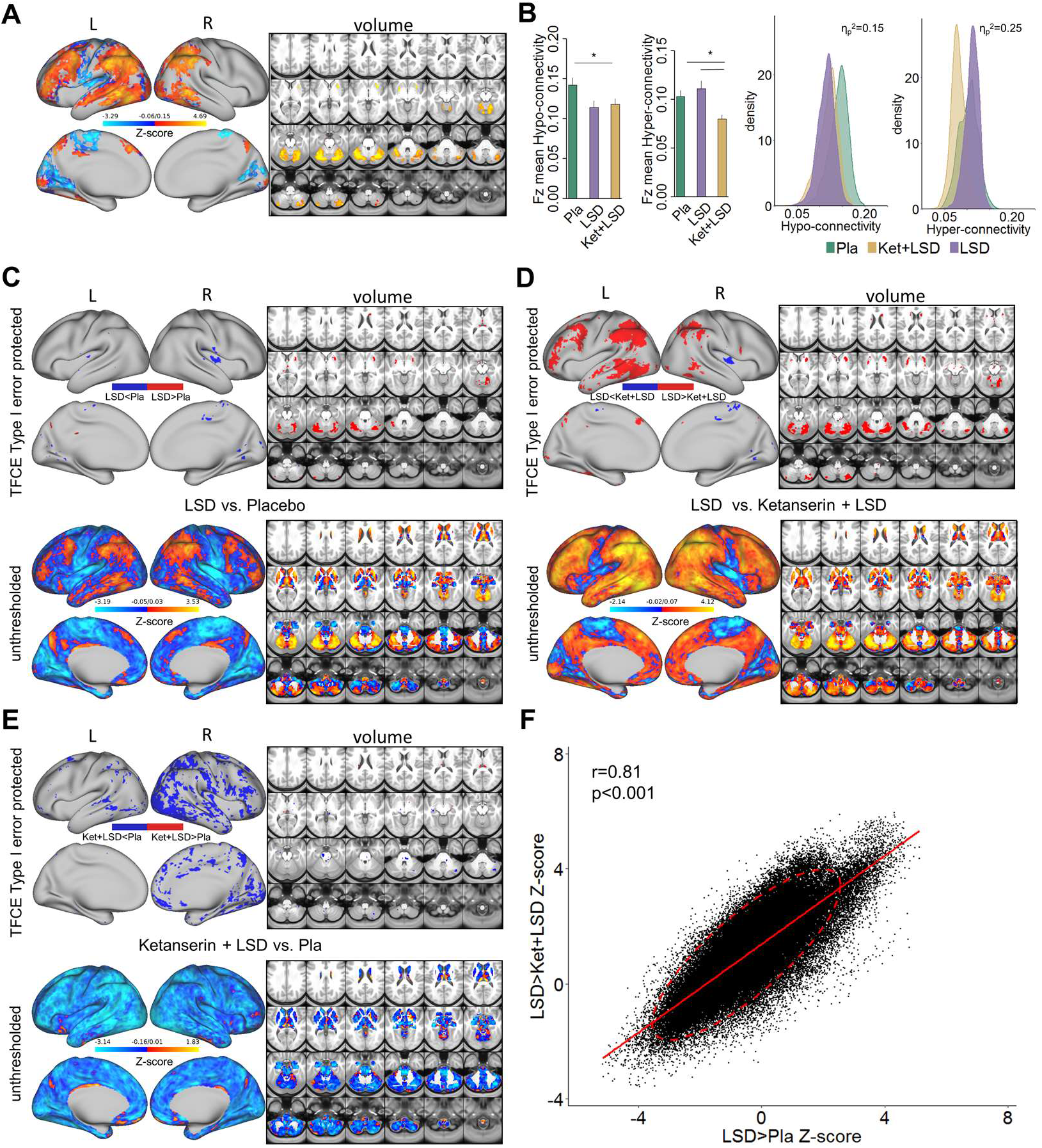
Effect of Drug Condition on GBC without GSR. **(A)** Z-score map for the effect of LSD condition vs. Ketanserin+LSD condition + Placebo condition within areas showing a significant main effect of drug (TFCE type 1 error protected). Red/orange areas indicate regions where participants exhibited stronger GBC in the LDS condition, whereas blue areas indicate regions where participants exhibited reduced GBC condition, compared with (Ketanserin+LSD)+Placebo conditions. **(B)** Bar plots show mean connectivity strength (Fz) values for hyper- and hypo-connected areas averaged across voxels showing a significant main effect of drug. Distribution plots show distribution of connectivity strength (Fz) values within voxels showing significant hyper- and hypo-connectivity for LSD compared to (Ketanserin+LSD)+Placebo conditions. **(C)** Top panel displays significant (TFCE type 1 error protected) areas showing increased (red) and decreased (blue) GBC in the LSD condition compared to Placebo. Lower panel shows the corresponding unthresholded Z-score map. Red/orange areas indicate regions where participants exhibited stronger GBC in the LSD condition, whereas blue areas indicate regions where participants exhibited reduced GBC in the LSD condition, compared with Placebo condition. **(D)** Top panel displays significant (TFCE type 1 error protected) areas showing increased (red) and decreased (blue) GBC in the LSD condition compared to Ketanserin+LSD. Lower panel shows the corresponding unthresholded Z-score map. Red/orange areas indicate regions where participants exhibited stronger GBC in the LSD condition, whereas blue areas indicate regions where participants exhibited reduced GBC in the LSD condition, compared with Ketanserin+LSD condition. **(E)** Top panel displays significant (TFCE type 1 error protected) areas showing increased (red) and decreased (blue) GBC in the Ketanserin+LSD condition compared to Placebo. Lower panel shows the corresponding unthresholded Z-score map. Red/orange areas indicate regions where participants exhibited stronger GBC in the Ketanserin+LSD condition, whereas blue areas indicate regions where participants exhibited reduced GBC in the Ketanserin+LSD condition, compared with Placebo condition. **(F)** Scatterplot showing a positive relationship between drug condition differences in GBC. Plotted are Z-scores for all voxels for the LSD>Placebo comparison (see panel C, X-axis) and LSD>Ketanserin+LSD comparison (see panel D, Y-axis). Ellipse marks the 95% confidence interval. N=24. * indicates significant difference between drug conditions, p<0.05, Bonferroni corrected.

**Figure S6.**
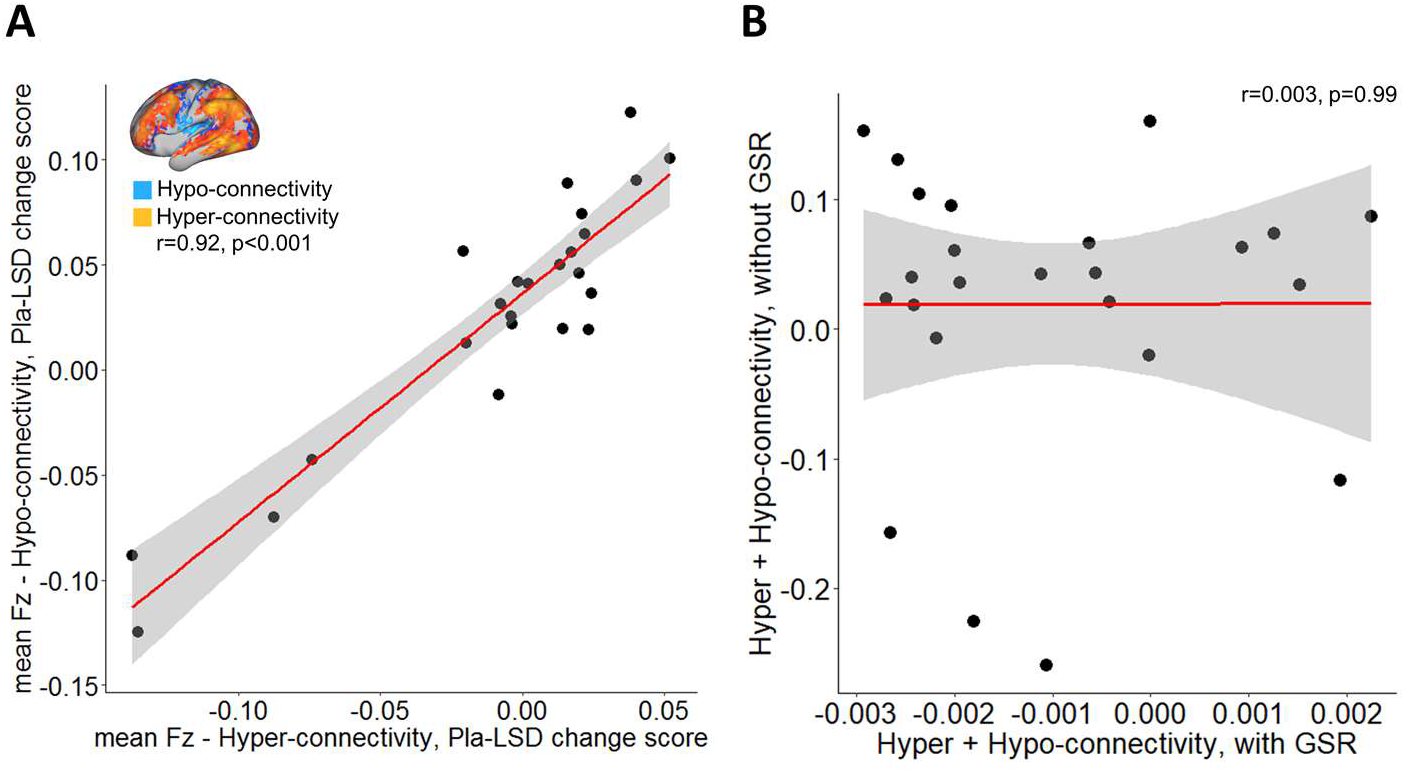
Relationship Between Hyper- and Hypo-connectivity Across Subjects. **(A)** Scatterplot showing significant positive relationship evident between averaged hyper- and hypo- connected voxels (based on the LSD vs (Ketanserin+LSD)+Placebo contrast without GSR, see inlet) across subjects (black data points) for placebo – LSD condition change scores. Grey background indicates the 95% confidence interval. **(B)** Scatterplot showing no significant relationship between Hyper + Hypo-connected Fz values with GSR (see Fig. 2) and Hyper + Hypo-connected Fz values without GSR (see **Fig S6A**). N=24. Grey shading indicates the 95% confidence interval.

**Figure S7.**
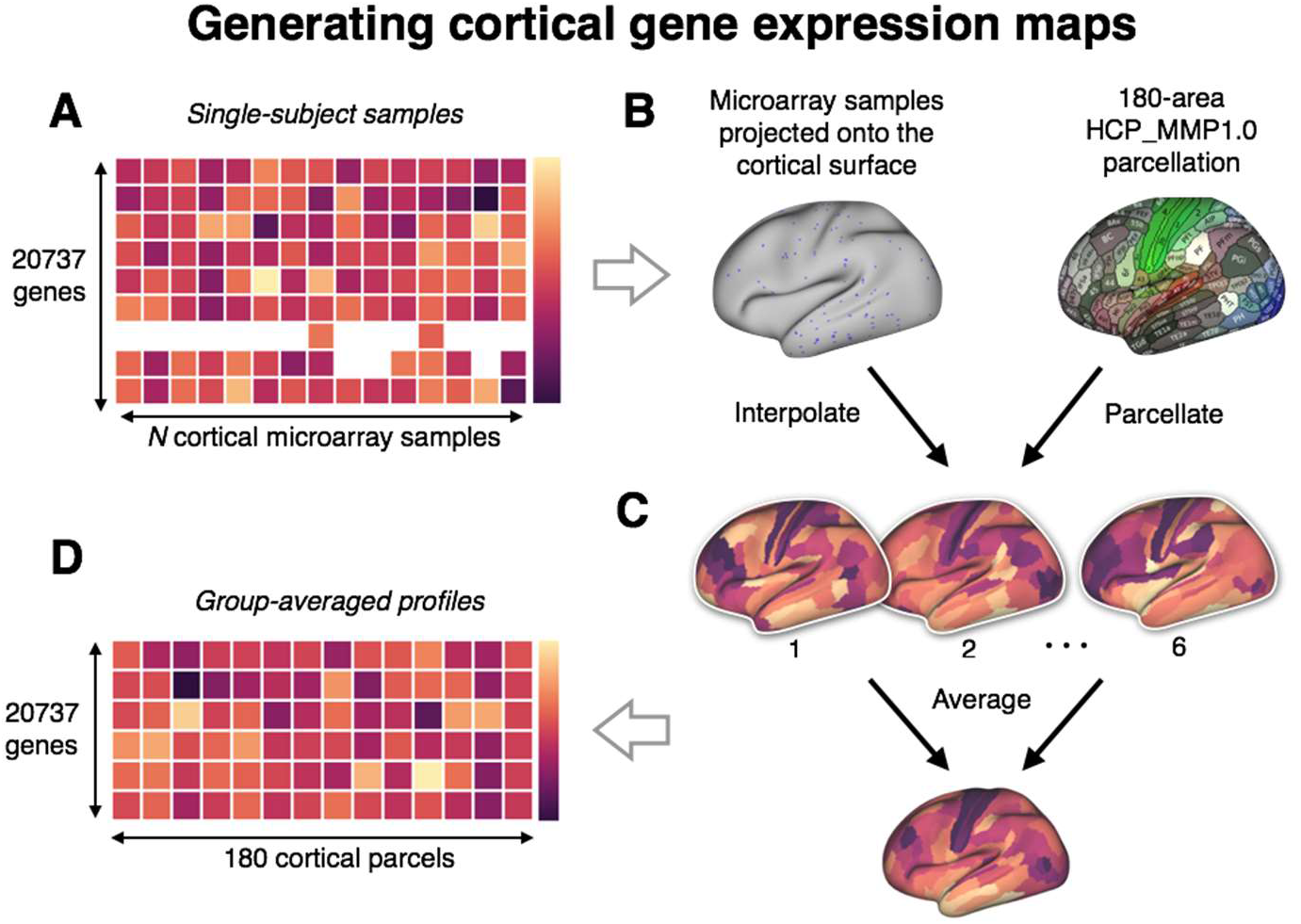
Schematic Illustrating the Process of Generating Cortical Gene Expression Maps from the Allen Human Brain Atlas (AHBA). **(A)** Using the AHBA (http://human.brain-map.org/), each subject’s microarray data is mapped onto the average cortical surface provided by the Human Connectome Project (HCP) **(B)** For each subject, the set of sparse gene expression sample locations on the cortical surface was used to generate a dense cortical map through a combination of interpolation via a Voronoi tessellation and parcellation using a 180-area multi-modal parcellaltion from the Human Connectome Project (HCP_MMP1.0) (20). **(C, D)** Single subject-level maps are averaged to construct the final group-averaged maps, which are used for all reported analyses. Complete details regarding the gene expression mapping protocol are reported in Burt et al.(21).

